# Latent space visualization, characterization, and generation of diverse vocal communication signals

**DOI:** 10.1101/870311

**Authors:** Tim Sainburg, Marvin Thielk, Timothy Q Gentner

## Abstract

Animals produce vocalizations that range in complexity from a single repeated call to hundreds of unique vocal elements patterned in sequences unfolding over hours. Characterizing complex vocalizations can require considerable effort and a deep intuition about each species’ vocal behavior. Even with a great deal of experience, human characterizations of animal communication can be affected by human perceptual biases. We present here a set of computational methods that center around projecting animal vocalizations into low dimensional latent representational spaces that are directly learned from data. We apply these methods to diverse datasets from over 20 species, including humans, bats, songbirds, mice, cetaceans, and nonhuman primates, enabling high-powered comparative analyses of unbiased acoustic features in the communicative repertoires across species. Latent projections uncover complex features of data in visually intuitive and quantifiable ways. We introduce methods for analyzing vocalizations as both discrete sequences and as continuous latent variables. Each method can be used to disentangle complex spectro-temporal structure and observe long-timescale organization in communication. Finally, we show how systematic sampling from latent representational spaces of vocalizations enables comprehensive investigations of perceptual and neural representations of complex and ecologically relevant acoustic feature spaces.

## 1 Author Summary

Of the thousands of species that communicate vocally, the repertoires of only a tiny minority have been characterized or studied in detail. This is due, in large part, to traditional analysis methods that require a high level of expertise that is hard to develop and often species-specific. Here, we present a set of novel methods to project animal vocalizations into latent feature spaces to quantitatively compare and develop visual intuitions about animal vocalizations, and to systematically synthesize novel species-typical vocalizations from learned feature sets. We demonstrate these methods across a series of analyses over 19 datasets of animal vocalizations from 29 different species, including songbirds, mice, monkeys, humans, and whales. We show how learned latent feature spaces untangle complex spectro-temporal structure, enable unbiased comparisons, and uncover high-level features such as individual identity and population dialects. We generate smoothly varying morphs between vocalizations from a songbird species with a spectro-temporally complex vocal repertoire, European starlings, and show how these methods enable a new degree of control over ecologically relevant signals that can be broadly applied across behavioral and physiological experimental settings.

## 2 Introduction

Vocal communication is a social behavior common to much of the animal kingdom in which acoustic signals are transmitted from sender to receiver to convey various forms of information such as identity, individual fitness, or the presence of danger. Across diverse fields, a set of pervasive research questions seeks to uncover the structure and mechanism of communication: What information is carried within signals? How are signals produced and received? How does the communicative transmission of information affect fitness and reproductive success? To approach these questions quantitatively, researchers rely largely on abstractions and characterizations of animal vocalizations [1]. For example, segmenting birdsong into discrete temporal elements and clustering these elements into discrete categories has played a crucial role in understanding syntactic structure in birdsong [1–9].

The characterization and abstraction of animal communicative signals remains both an art and a science. For example, Kershenbaum et. al., [1] survey and describe the most common analysis pipeline used to abstract and describe vocal sequences and find that analyses are broadly comprised of the same pattern of steps: (1) the collection of data, (2) segmentation of vocalizations into units, (3) characterization of sequences, and (4) identification of meaning. Within this more general paradigm however, a number of heuristics exist for determining how best to segment, label, and characterize vocalizations. It remains largely up to experimenter expertise to determine which heuristics to apply. For instance, how do we determine what constitutes a ‘unit’ of humpback whale song? Communicative repertoires of different species vary widely, and their characterization can be difficult, time-consuming, and can often rely heavily upon deep intuitions about the structure of vocal repertoires formed by experts in a species’ communication. When such intuitions are available they should be considered, of course, but they are generally rare in comparison to the wide range of communication signals observed. Thus, communication remains understudied in most of the thousands of vocally communicating species. Even in well-documented model species for vocal communication, characterizations of vocalizations are often influenced by human-centric biases and heuristics [1, 10–12]. We therefore turn to unsupervised computational methodology to aid in the role of characterizing vocal communication. One area where machine learning has flourished is the representation of complex statistical patterns in data. In many different domains, machine learning methods have uncovered and untangled meaningful representations of data based upon the statistics of their structure [13, 14, 14–16, 16, 17]. In the characterization of animal communication, these techniques are therefore well suited to quantitatively investigate complex statistical structure present in vocal data that otherwise rely upon expert intuitions. In this paper, we demonstrate the utility of unsupervised latent models, statistical models that learn latent representations of complex vocal data, in describing animal communication.

### 2.1 Latent models of acoustic communication

The utility of the latent models we describe here can be broadly divided into two categories: dimensionality reduction, and generativity. Dimensionality reduction refers to the projection of high-dimensional data, such as syllables of birdsong, into a smaller number of dimensions, while retaining the structure and variance present in the original high-dimensional data. Each point in that high-dimensional space (e.g. a syllable of birdsong) can be projected into that lower-dimensional ‘latent’ feature space and each dimension can be thought of as a feature of the dataset. Generativity refers to the process of sampling from the low-dimensional latent space and generating novel data in the original high dimensional space.

The traditional practice of developing a set of basis-features on which vocalizations can be quantitatively compared is a form of dimensionality reduction and comes standard in most animal vocalization analysis software (e.g. Luscinia [18], Sound Analysis Pro [19, 20], Avisoft [21], and Raven [22]). Birdsong, for example, is often analyzed on the basis of features such as amplitude envelope, Weiner entropy, spectral continuity, pitch, duration, and frequency modulation [1, 19]. Likewise, grouping elements of animal vocalizations (e.g. syllables of birdsong, mouse ultrasonic vocalizations) into abstracted discrete categories can be thought of as dimensionality reduction, where each category is a single orthogonal dimension. In machine learning parlance, the process of determining the relevant features, or dimensions, of a particular dataset, is called *feature engineering*. Engineered features are ideal for many analyses because they are human-interpretable in models that describe the relative contribution of those features as explanatory variables, for example explaining the contribution of the fundamental frequency of a *coo* call in predicting caller identity in macaques [23]. Feature engineering, however, has two caveats. First, the features selected by humans are biased by human perceptual systems, which are not necessarily “tuned” for analyzing all non-human communication signals. Second, feature engineering typically requires significant domain knowledge, which is time-consuming to acquire and difficult to generalize across species, impairing cross-species comparisons.

An attractive alternative approach is to project animal vocalizations into low-dimensional feature spaces that are determined directly from the structure of the data. For example, animal vocalizations can be projected into linear feature spaces using principal component analysis where each successive dimension represents an orthogonal transformation capturing the maximal variance possible in the data [1, 24], or vocalizations can be decomposed into features using linear discriminant analysis where features are determined by their ability to explain variance in a specific dimension of the data, such as individual identity [25]. Dimensionality reduction can also be nonlinear, allowing for representations that better capture relationships between data (e.g. the similarity between two syllables of birdsong). The utility of non-linear dimensionality reduction techniques are just now coming to fruition in the study of animal communication, for example using t-distributed stochastic neighborhood embedding (t-SNE; [26]) to describe the development of zebra finch song [27], using Uniform Manifold Approximation and Projection (UMAP; [28]) to describe and infer categories in birdsong [3, 29], or using deep neural networks to synthesize naturalistic acoustic stimuli [30, 31]. Developments in non-linear representation learning have helped fuel the most recent advancements in machine learning, untangling statistical relationships in ways that provide more explanatory power over data than traditional linear techniques [13, 14]. These advances have proven important for understanding data in diverse fields including the life sciences (e.g. [3, 16, 27, 29, 32, 33]).

In this paper, we propose and give a broad overview of latent models that learn complex feature-spaces of vocalizations, requiring few *a priori* assumptions about a species’ vocalizations. We utilize UMAP [28] along with several generative neural networks [34–37] for data representation and generation. UMAP is a recent method for non-linear dimensionality reduction that, grounded firmly in category theory, projects data into a lower-dimensional space while preserving as much of the local and global structure of the data manifold as possible. We chose to use UMAP over t-SNE, a related and longer-standing dimensionality reduction algorithm, because UMAP has been shown to preserve more global structure, decrease computation time, and effectively produce more meaningful data representations in a number of areas within the natural sciences (e.g. [3, 16, 28, 29]). We show that these methods reveal informative feature-spaces that enable the formulation and testing of hypotheses about animal communication. In addition, these methods allow for systematic sampling from complex feature spaces of animal communicative signals, providing a high degree of control control over vocal signals in real-world experiments. We apply our method to diverse datasets consisting of over 20 species (Table 2), including humans, bats, songbirds, mice, cetaceans, and nonhuman primates. We introduce methods for treating vocalizations both as sequences of temporally discrete elements such as syllables, as is traditional in studying animal communication [1], as well as temporally continuous trajectories, as is becoming increasingly common in representing neural sequences [38]. Using both methods, we show that latent projections produce visually-intuitive and quantifiable representations that capture complex acoustic features. We show comparatively that the spectrotemporal characteristics of vocal units vary from species to species in how distributionally discrete they are and discuss the relative utility of different ways to represent different communicative signals. Finally, we show an example of how latent models allow animal vocal repertoires to be systematically exploited to probe the perceptual and neural representations of vocal signals without degrading their complex underlying spectrotemporal structure.

## 3 Results

### 3.1 Discrete latent projections of animal vocalizations

To explore the broad utility of latent models in capturing features of vocal repertoires, we analyzed nineteen datasets consisting of 400 hours of vocalizations and over 3,000,000 discrete vocal units from 29 unique species (Table 2). Each vocalization dataset was temporally segmented into discrete units (e.g. syllables, notes), either based upon segmentation boundaries provided by the dataset (where available), or using a novel dynamic-thresholding segmentation algorithm that segments syllables of vocalizations between detected pauses in the vocal stream (Fig 16; See Segmentation). Each dataset was chosen because it contains large repertoires of vocalizations from relatively acoustically isolated individuals that can be cleanly separated into temporally-discrete vocal units. With each temporally discrete vocal unit we computed a spectrographic representation (Supplementary Fig 17; See Spectrogramming). We then projected the spectrographic representation into latent feature spaces using UMAP (Figs 1, 2, 3, 4). From these latent feature spaces, we analyzed datasets for several features, including stereotypy/clusterability (Figs 1, 5), individual identity (Fig 2), species identity (Fig 3A,B), geographic populations (Fig 3C), speech features (Figs 4, 18), and sequential organization (Fig 6).

**Figure 1:**
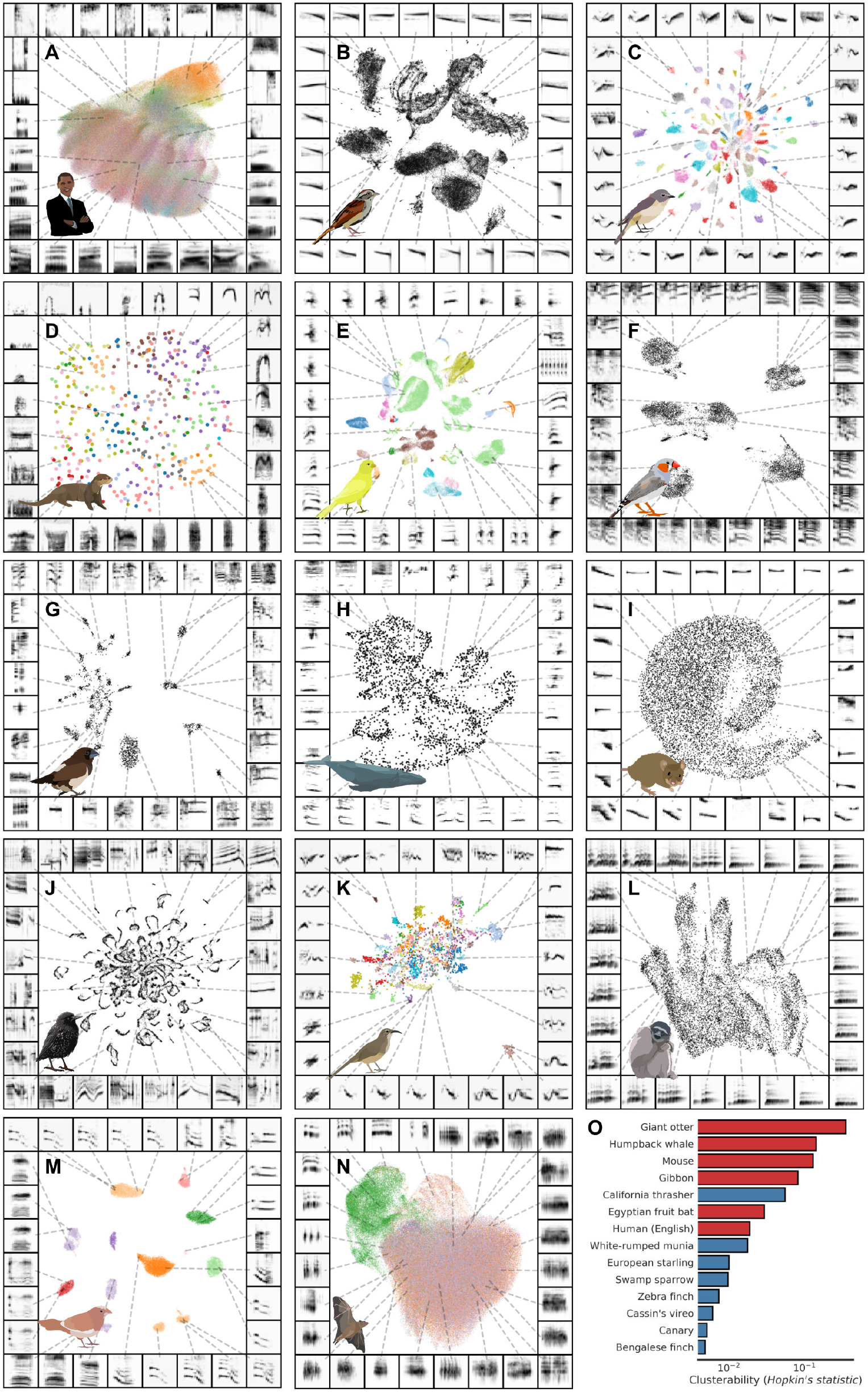
UMAP projections of vocal repertoires across diverse species. Each plot shows vocal elements discretized, spectrogrammed, and then embedded into a 2D UMAP space, where each point in the scatterplot represents a single element (e.g. syllable of birdsong). Scatterplots are colored by element categories where available. The borders around each plot are example spectrograms pointing toward different regions of the scatterplot. Plots are shown for single individuals in datasets where vocal repertoires were visually observed to be distinct across individuals (E, F, G, J, M), and are shown across individuals for the remainder of plots. (A) Human phonemes. (B) Swamp sparrow notes. (C) Cassin’s vireo syllables. (D) Giant otter calls. (E) Canary syllables. (F) Zebra finch sub-motif syllables. (G) White-rumped munia syllables. (H) Humpback whale syllables. (I) Mouse USVs. (J) European starling syllables. (K) California thrasher syllables. (L) Gibbon syllables. (M) Bengalese finch syllables. (N) Egyptian fruit bat calls (color is context). (O) Clusterability (Hopkin’s metric) for each dataset. Lower is more clusterable. Hopkin’s metric is computed over UMAP projected vocalizations for each species. Color represents species category (red: mammal, blue: songbird).

**Figure 2:**
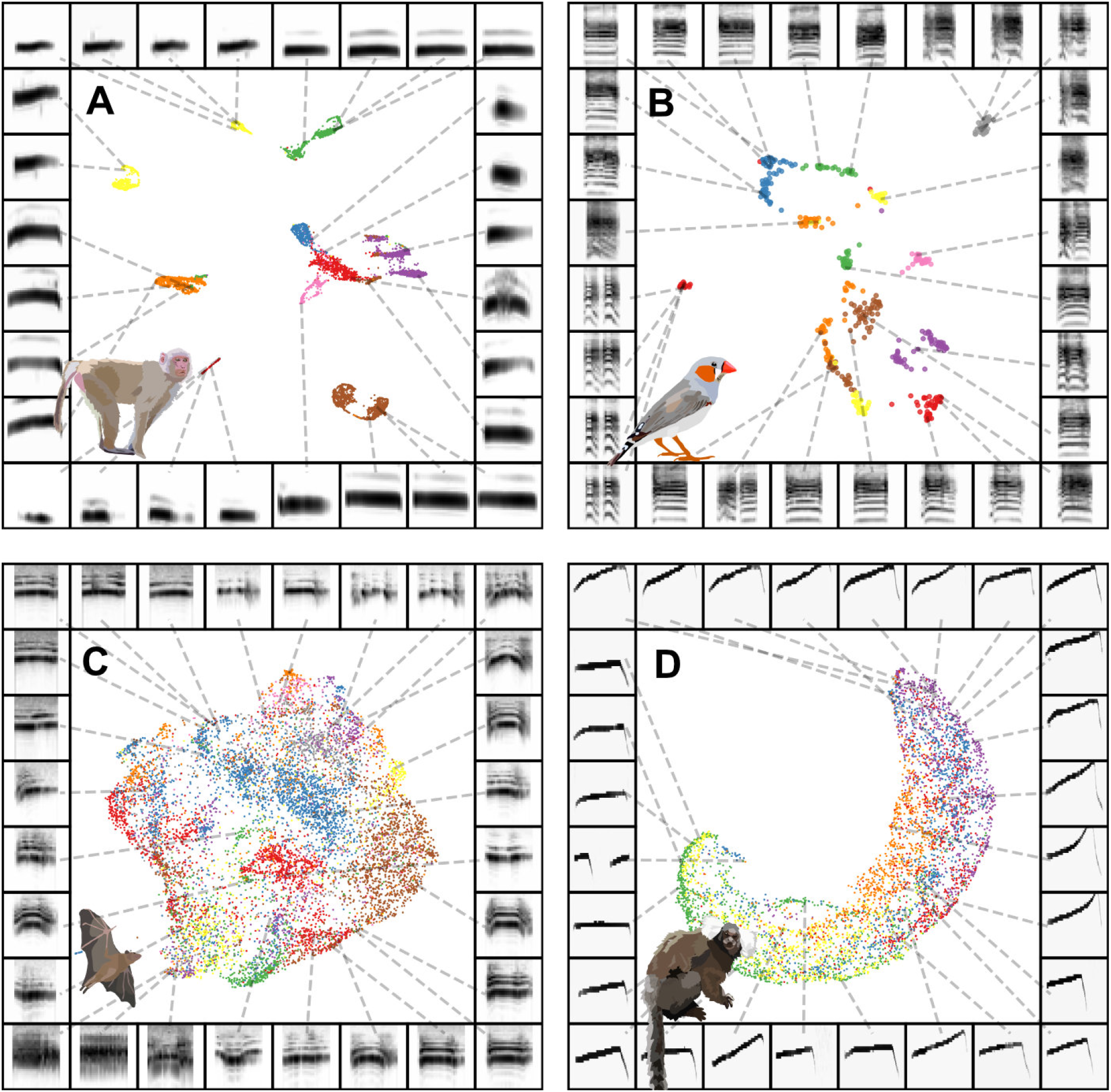
Individual identity is captured in projections for some datasets. Each plot shows vocal elements discretized, spectrogrammed, and then embedded into a 2D UMAP space, where each point in the scatterplot represents a single element (e.g. syllable of birdsong). Scatterplots are colored by individual identity. The borders around each plot are example spectrograms pointing toward different regions of the scatterplot. (A) Rhesus macaque coo calls. (B) Zebra finch distance calls. (C) Fruit bat infant isolation calls. (D) Marmoset phee calls.

**Figure 3:**
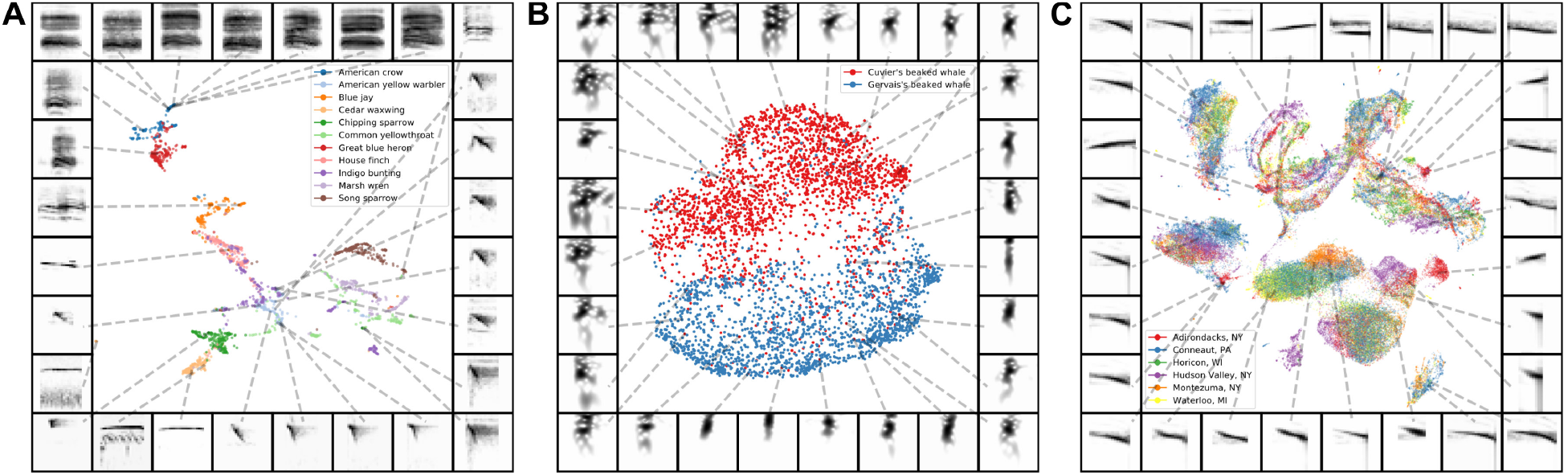
Comparing species with latent projections. (A) Calls from eleven species of North American birds are projected into the same UMAP latent space. (B) Cuvier’s and Gervais’s beaked whale echolocation clicks are projected into UMAP latent space and fall into two discrete clusters. (C) Notes of swamp sparrow song from six different geographical populations.

**Figure 4:**
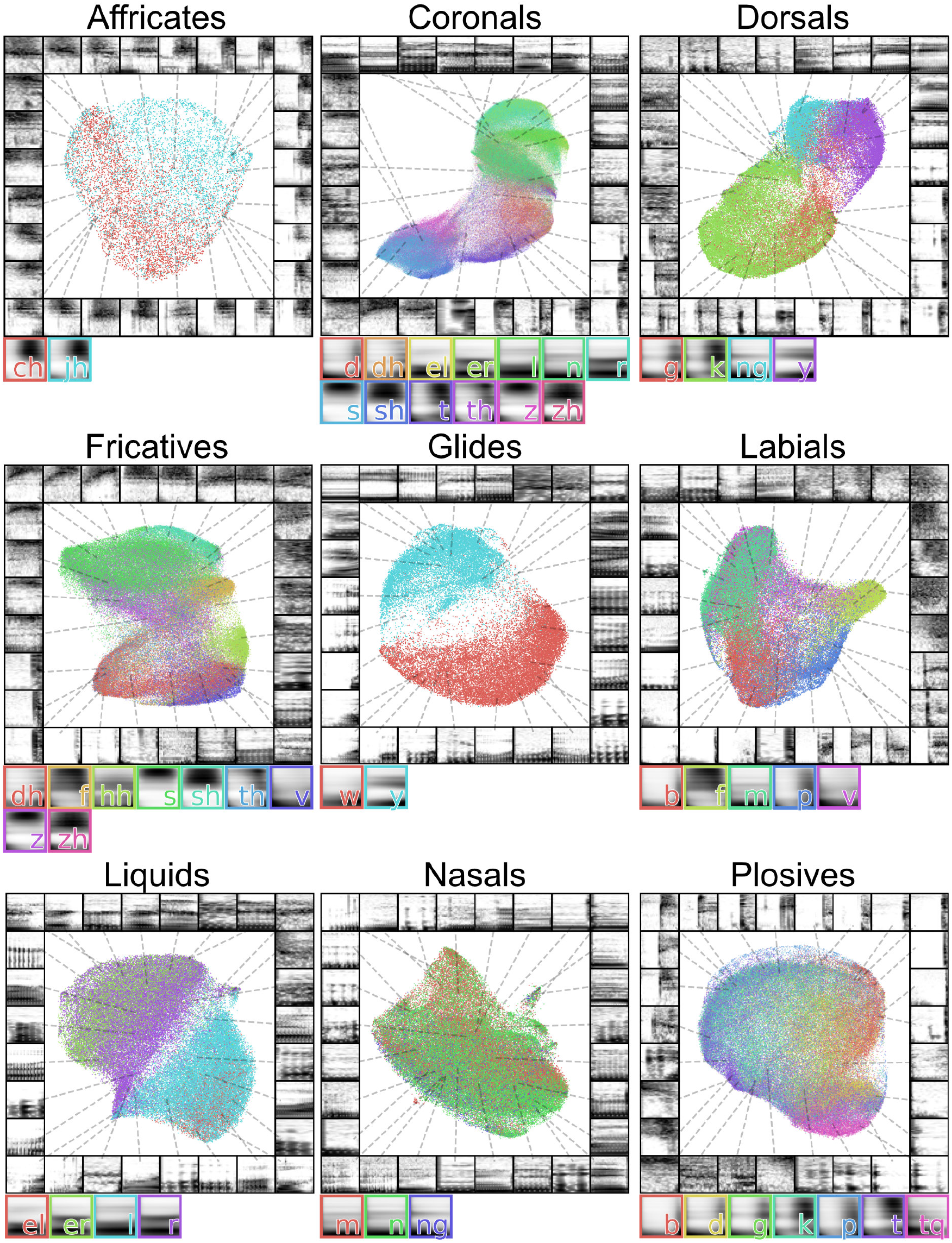
Latent projections of consonants. Each plot shows a different set of consonants grouped by phonetic features. The average spectrogram for each consonant is shown to the right of each plot.

**Figure 5:**
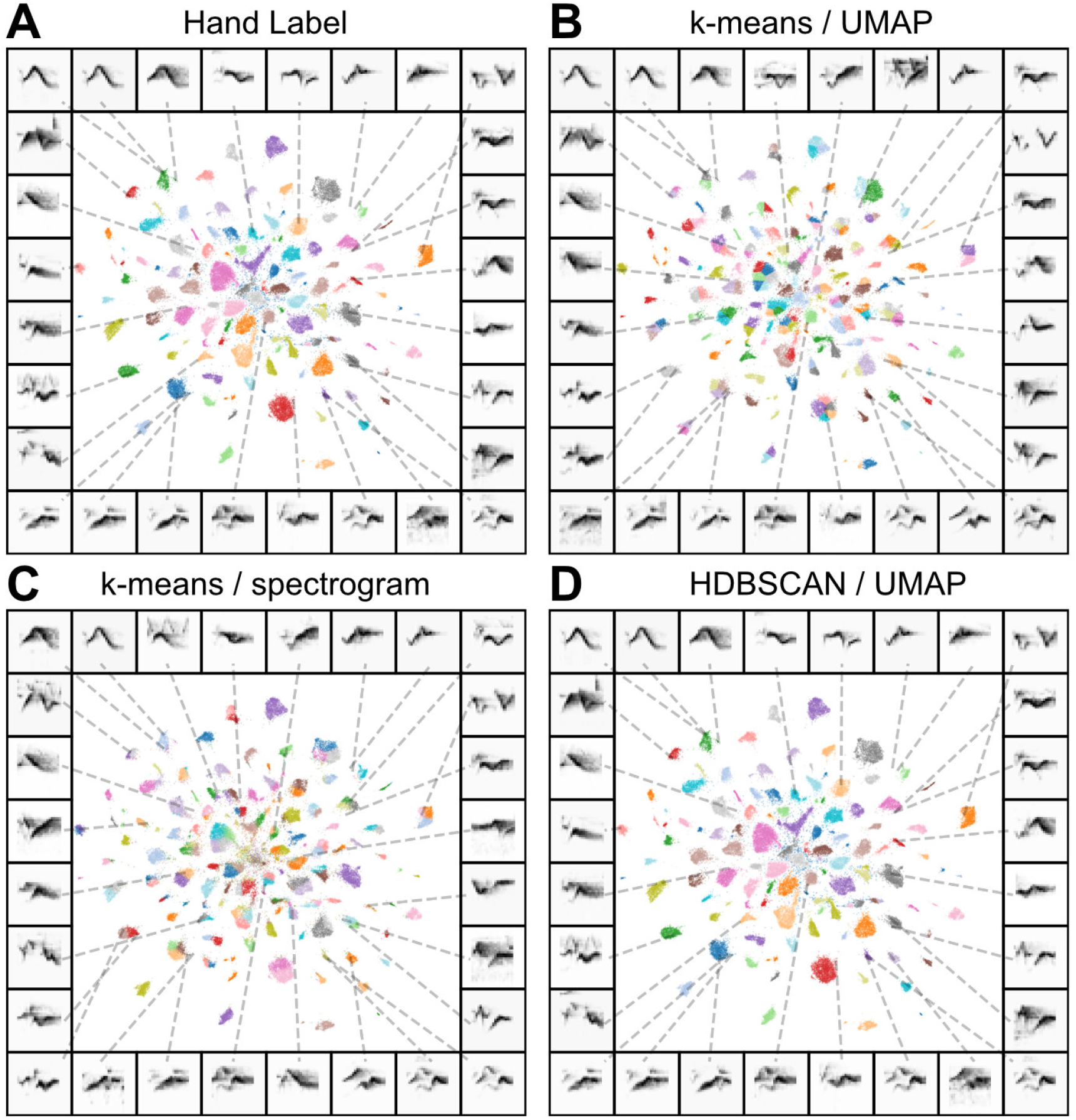
Cassin’s vireo syllables projected into UMAP and clustered algorithmically. (A) A scatterplot where each syllable is a single point projected into two UMAP dimensions. Points are colored by their hand-labeled categories, which generally fall into discrete clusters in UMAP space. Each other frame is the same scatterplot, where colors are cluster labels produced using (B) k-means over UMAP projections (C) k-means directly on syllable spectrograms (D) HDBSCAN on UMAP projections.

**Figure 6:**
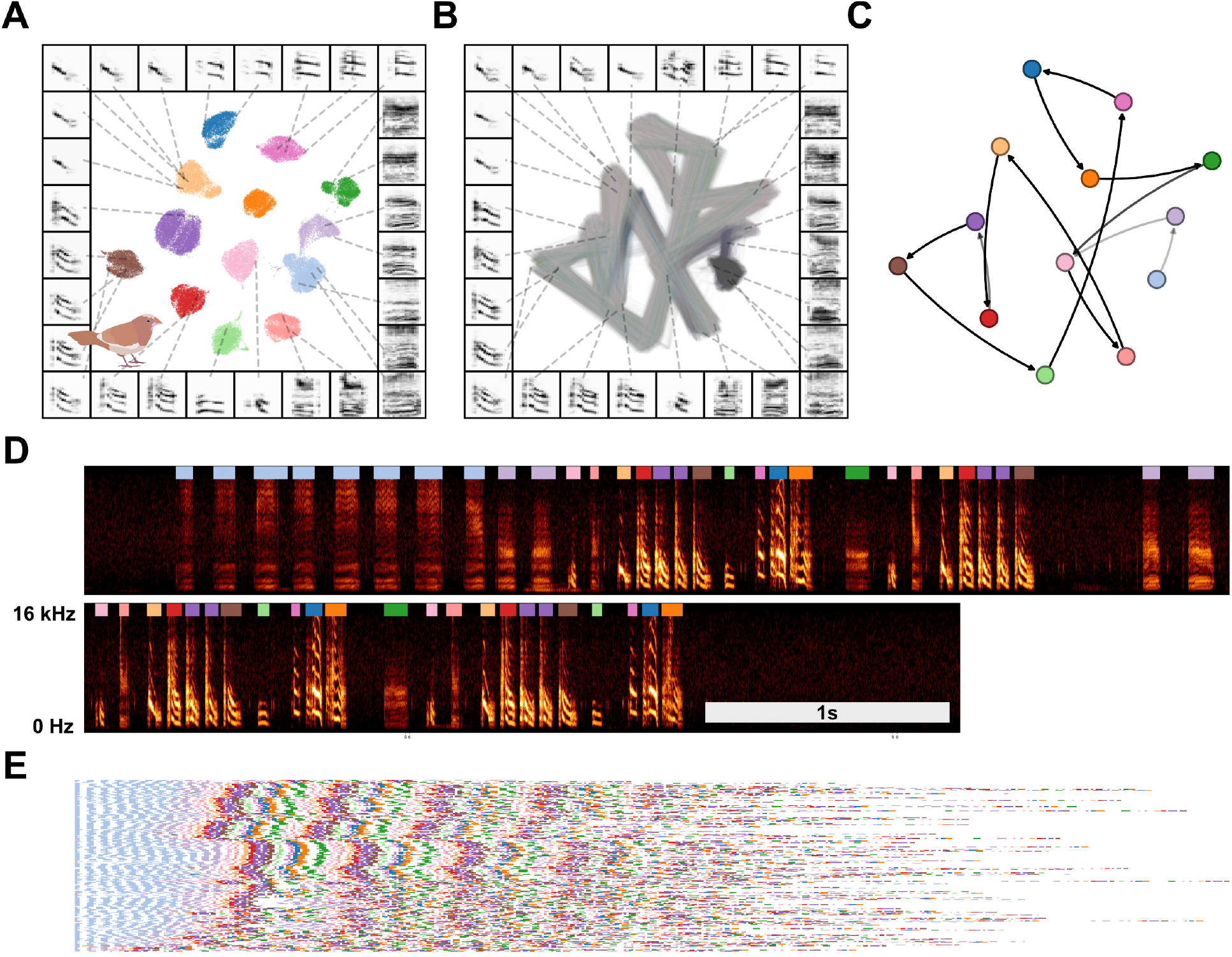
Latent visualizations of Bengalese finch song sequences. (A) Syllables of Bengalese finch songs from one individual are projected into 2D UMAP latent space and clustered using HDBSCAN. (B) Transitions between elements of song are visualized as line segments, where the color of the line segment represents its position within a bout. (C) The syllable categories and transitions in (A) and (B) can be abstracted to transition probabilities between syllable categories, as in a Markov model. (D) An example vocalization from the same individual, with syllable clusters from (A) shown above each syllable. (E) A series of song bouts. Each row is one bout, showing overlapping structure in syllable sequences. Bouts are sorted by similarity to help show structure in song.

#### 3.1.1 Variation in discrete distributions and stereotypy

In specices as phylogenetically diverse as songbirds and rock hyraxes, analyzing the sequential organization of communication relies upon similar methods of segmentation and categorization of discrete vocal elements [1]. In species such as the Bengalese finch, where syllables are highly stereotyped, clustering syllables into discrete categories is a natural way to abstract song. The utility of clustering song elements in other species, however, is more contentious because discrete category boundaries are not as easily discerned [10, 11, 29, 39]. We looked at how discrete the clusters found in UMAP latent spaces are across vocal repertoires from different species.

Visually inspecting the latent projections of vocalizations (Fig 1) reveals appreciable variability in how the repertoires of different species cluster in latent space. For example, mouse USVs appear as a single cluster (Fig 1I), while finch syllables appear as multiple discrete clusters (Fig 1M,F), and gibbon song sits somewhere in between (Fig 1L). This suggests that the spectro-temporal acoustic diversity of vocal repertoires fall along a continuum ranging from unclustered, uni-modal to highly clustered.

We quantified the varying clusterability of vocal elements in each species by computing the Hopkin’s statistic (Eq. 3; See Clusterability section) over latent projections for each dataset in Fig 1. The Hopkin’s statistic captures how far a distribution deviates from uniform random and give a common measure of the ‘clusterability’ of a dataset [40]. There is a clear divide in clusterability between mammalian and songbird repertoires, where the elements of songbird repertoires tend to cluster more than mammalian vocal elements (Fig 1O). The stereotypy of songbird (and other avian) vocal elements is well documented [41, 42] and at least in zebra finches is related to the high temporal precision in the singing-related neural activity of vocal-motor brain regions [43–45].

#### 3.1.2 Vocal features

Latent non-linear projections often untangle complex features of data in human-interpretable ways. For example, the latent spaces of some neural networks linearize the presence of a beard in an image of a face without being trained on beards in any explicit way [15, 35]. Complex features of vocalizations are similarly captured in intuitive ways in latent projections [3, 29–31]. Depending on the organization of the dataset projected into a latent space, these features can extend over biologically or psychologically relevant scales. Accordingly, we used our latent models to look at spectro-temporal structure within the vocal repertoires of individual’s, and across individuals, populations, and phylogeny. These latent projections capture a range of complex features, including individual identity (Fig 2), species identity (Fig 3A,B), linguistic features (Figs 4, 18), syllabic categories (Figs 6, 5, 1, 7), and geographical distribution (Fig 3C). We discuss each of these complex features in more detail below.

**Figure 7:**
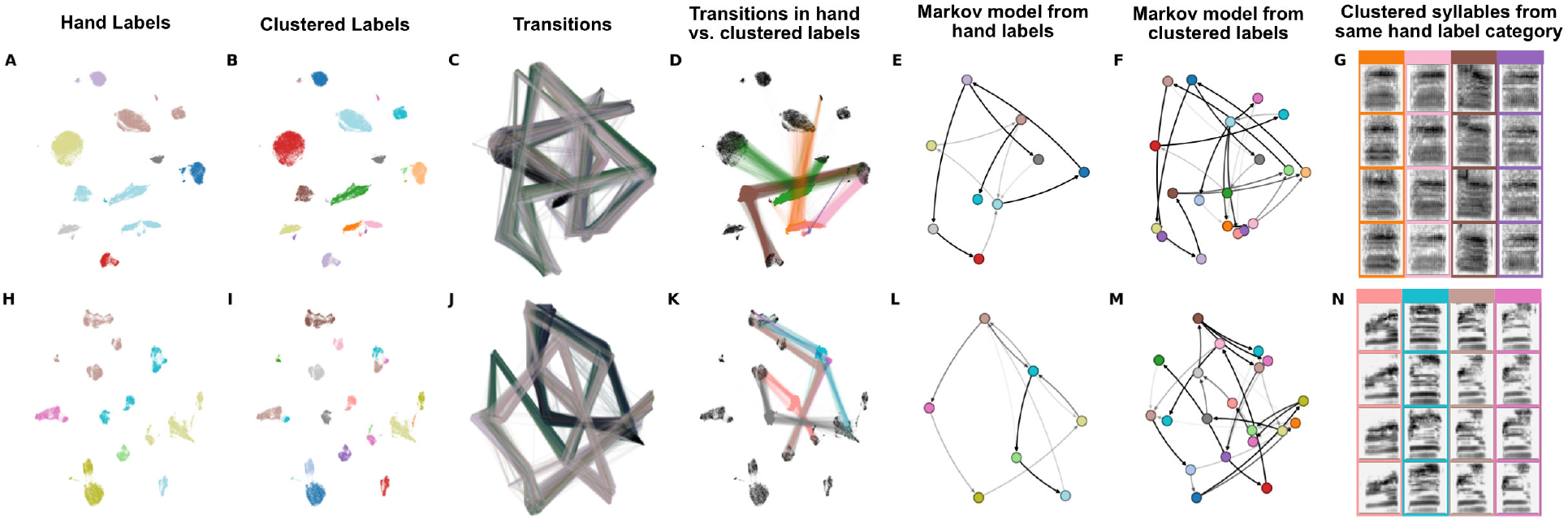
Latent comparisons of hand- and algorithmically-clustered Bengalese finch song. A-G are from a dataset produced by Nicholson et al., [9] and H-N are from a dataset produced by Koumura et al., [10] (A,H) UMAP projections of syllables of Bengalese finch song, colored by hand labels. (B,I) Algorithmic labels (UMAP/HDBSCAN). (C, J) Transitions between syllables, where color represents time within a bout of song. (D,K) Comparing the transitions between elements from a single hand-labeled category that comprises multiple algorithmically labeled clusters. Each algorithmically labeled cluster and the corresponding incoming and outgoing transitions are colored. Transitions to different regions of the UMAP projections demonstrate that the algorithmic clustering method finds clusters with different syntactic roles within hand-labeled categories. (E,L) Markov model from hand labels colored the same as in (A,H) (F,M) Markov model from clustered labels, colored the same as in (B,I). (G,H) Examples of syllables from multiple algorithmic clusters falling under a single hand-labeled cluster. Colored bounding boxes around each syllable denotes the color category from (D,K).

##### Individual identity

Many species produce caller-specific vocalizations that facilitate the identification of individuals when other sensory cues, such as sight, are not available. The features of vocalizations facilitating individual identification vary, however, between species. We projected identity call datasets (i.e., sets of calls thought to carry individual identity information) from four different species into UMAP latent spaces (one per species) to observe whether individual identity falls out naturally within the latent space.

We looked at four datasets where both caller and call-type are available. Caller identity is evident in latent projections of all four datasets (Fig 2). The first dataset is comprised of Macaque coo calls, where identity information is thought to be distributed across multiple features including fundamental frequency, duration, and Weiner entropy [23]. Indeed, the latent projection of coo calls clustered tightly by individual identity (Fig 2A). The same is true for Zebra finch distance calls (Fig 2B), Egyptian fruit bat pup isolation calls (Fig 2C), which in other bat species are discriminable by adult females [46, 46, 47], and Marmoset phee calls. It is perhaps interesting, given the range of potential features thought to carry individual identity [23] that the phee calls appear to lie along a single continuum (Fig 2D). These patterns suggest that some calls, such as macaque *coo* calls, are likely more differentiable than other calls, such as marmoset *phee* calls.

These latent projections demonstrate that caller identity can be obtained from all these vocalizations, and, we note, without *a priori* knowledge of specific spectro-temporal features. Because no caller identity information is used in learning the latent projections, the emergence of this information indicates that the similarity of within-caller vocalizations contains enough statistical power to overcome variability between callers. This within-caller structure likely facilitates conspecific learning of individual identity without *a priori* expectations for the distribution of relevant features [48], in the same way that developing sensory systems adapt to natural environmental statistics [49].

##### Cross species comparisons

Classical comparative studies of vocalizations across species rely on experience with multiple species’ vocal repertoires. This constrains comparisons to those species whose vocalizations are understood in similar features spaces, or forces the choice of common feature spaces that may obscure relevant variation differently in different species. Because latent models learn arbitrary complex features of datasets, they can yield less biased comparisons between vocal repertoires where the relevant axes are unknown, and where the surface structures are either very different, for example canary and starling song, or very similar, like the echolocation clicks of two closely related beaked whales.

To explore how well latent projections capture vocal repertoire variation across species, we projected a dataset containing monosyllabic vocalizations [50] from eleven different species of North American birds into UMAP latent space. Similar “calls”, like those from the American crow *caw* and great blue heron *roh* are closer together in latent space, while more distinct vocalizations, like chipping sparrow notes, are further apart (Fig 3A). Latent projections like this have the potential power to enable comparisons across broad phylogenies without requiring decisions about which acoustic features to compare.

At the other extreme is the common challenge in bioacoustics research to differentiate between species with very similar vocal repertoires. For example, Cuvier’s and Gervais’ beaked whales, two sympatric species recorded in the Gulf of Mexico, have echolocation clicks with highly overlapping power spectra that are generally differentiated using supervised learning approaches (c.f. [51, 52]). We projected a dataset containing Cuvier’s and Gervais’ beaked whale echolocation clicks into UMAP latent space. Species-identity again falls out nicely, with clicks assorting into distinct clusters that correspond to each species (Fig 3B).

##### Population geography

Some vocal learning species produce different vocal repertoires (dialects) across populations. Differences in regional dialects across populations are borne out in the categorical perception of notes [53–55], much the same as cross-linguistic differences in the categorical perception of phonemes in human speech [56]. To compared vocalizations across geographical populations in the swamp sparrow, which produces regional dialects in its trill-like songs [18], we projected individual notes into a UMAP latent space. Although the macro-structure of clusters suggest common note-types for the entire species, most of the larger clusters show multiple clear sub-regions that are tied to vocal differences between geographical populations (Fig 3C).

##### Phonological features

The sound segments that make up spoken human language can be described by distinctive phonological features that are grouped according to articulation place and manner, glottal state, and vowel space. A natural way to look more closely at variation in phoneme production is to look at variation between phonemes that comprise the same phonological features. As an example, we projected sets of consonants that shared individual phonological features into UMAP latent space (Figs 4, 18). In most cases, individual phonemes tended to project to distinct regions of latent space based upon phonetic category, and consistent with their perceptual categorization. At the same time, we note that latent projections vary smoothly from one category to the next, rather than falling into discrete clusters. This provides a framework that could be used in future work to characterize the distributional properties of speech sounds in an unbiased manner. Likewise, it would be interesting to contrast projections of phonemes from multiple languages, in a similar manner as the swamp sparrow (Fig 3C), to visualize and characterize variation in phonetic categories across languages [56].

#### 3.1.3 Clustering vocal element categories

Unlike human speech, UMAP projections of birdsongs fall more neatly into discriminable clusters (Fig 1). If clusters in latent space are highly similar to experimenter-labeled element categories, unsupervised latent clustering could provide an automated and less time-intensive alternative to hand-labeling elements of vocalizations. To examine this, we compared how well clusters in latent space correspond to experimenter-labeled categories in three human-labeled datasets: two separate Bengalese finch datasets [57, 58], and one Cassin’s vireo dataset [7]. We compared three different labeling techniques: a hierarchical density-based clustering algorithm (HDBSCAN; [59]) applied over latent projections in UMAP, k-means [60] clustering applied over UMAP, and k-means clustering applied over spectrograms (Fig 5; Table 1). To make the k-means algorithm more competitive with HDBSCAN, we set the number of clusters in k-means equal to the number of clusters in the hand-clustered dataset, while HDBSCAN was not parameterized at all. We computed the similarity between hand and algorithmically labeled datasets using four different metrics (See Methods section). For all three datasets, HDBSCAN clustering over UMAP projections is most similar to hand labels and visually overlaps best with clusters in latent space (Fig 5; Table 1). These results show that latent projections facilitate unsupervised clustering of vocal elements into human-like syllable categories better than spectrographic representations alone. At the same time, latent clusters do not always exactly match experimenter labels, a phenomenon that we explore in greater depth in the next section.

**Table 1:** Cluster similarity to ground truth labels for two Bengalese finch and one Cassin’s vireo dataset. Three clustering methods were used: (1) HDBSCAN clustering of UMAP projections (2) KMeans on spectrograms (3) KMeans on UMAP projections. KMeans was initialized with the correct number of clusters to make it more competitive with HDBSCAN clustering.

|  |  | Homogeneity | Completeness | V-Measure | Adjusted MI |
| --- | --- | --- | --- | --- | --- |
| **B. Finch (Kourmura)** |  |  |  |  |  |
|  | **HDBSCAN/UMAP** | 0.992±0.005 | 0.849±0.091 | 0.912±0.054 | 0.849±0.091 |
|  | **KMeans** | 0.903±0.035 | 0.836±0.068 | 0.867±0.047 | 0.834±0.065 |
|  | **KMeans/UMAP** | 0.904±0.078 | 0.855±0.095 | 0.878±0.084 | 0.852±0.092 |
| **B. Finch (Nicholson)** |  |  |  |  |  |
|  | **HDBSCAN/UMAP** | 0.94±0.052 | 0.862±0.081 | 0.895±0.033 | 0.832±0.049 |
|  | **KMeans** | 0.957±0.022 | 0.699±0.1 | 0.805±0.074 | 0.699±0.1 |
|  | **KMeans/UMAP** | 0.96±0.026 | 0.675±0.103 | 0.789±0.08 | 0.674±0.103 |
| **Cassin's vireo** |  |  |  |  |  |
|  | **HDBSCAN/UMAP** | 0.934 | 0.935 | 0.934 | 0.932 |
|  | **KMeans** | 0.895 | 0.808 | 0.849 | 0.801 |
|  | **KMeans/UMAP** | 0.926 | 0.827 | 0.874 | 0.821 |

**Table 2:** Overview of the species and datasets used in this paper

| Species | # Indv. | # Elements | Median len. (s) | Total length (s) | # Rec. | References |
| --- | --- | --- | --- | --- | --- | --- |
| American crow | Unk. | syllables: 252 | syllables: 0.37 | 100.5 | 252 | [50, 108] |
| Bengalese finch | 4 | syllables: 215480 | syllables: 0.065 | 40205.6 | 2663 | [57] |
| Bengalese finch | 11 | notes: 214915 | notes: 0.089 | 35365.9 | 2964 | [8, 58] |
| Blue jay | Unk. | syllables: 250 | syllables: 0.47 | 141.2 | 250 | [50, 108] |
| California thrasher | 18 | syllables: 15328 | syllables: 0.146 | 19958.9 | 92 | [6, 96] |
| Canary | 5 | phrases: 22167 | phrases: 1.319 | 36986.9 | 2320 | [5] |
|  |  | syllables: 497338 | syllables: 0.04 |  |  |  |
| Cassin's vireo | 48 | syllables: 67316 | syllables: 0.332 | 434782.4 | 422 | [7, 96] |
| Cedar waxwind | Unk. | syllables: 245 | syllables: 0.425 | 116.0 | 245 | [50, 108] |
| Chipping sparrow | Unk. | syllables: 252 | syllables: 0.09 | 24.9 | 252 | [50, 108] |
| Common marmoset | 33 | calls: 14289 | calls: 1.084 | 76400.7 | 768 | [25] |
| Common yellowthroat | Unk. | syllables: 255 | syllables: 0.1 | 35.4 | 255 | [50, 108] |
| Cuvier's beaked whale | Unk. | clicks: 2237 | clicks: 0.001 | 2.3 | 2237 | [51, 107] |
| Egyptian fruit bat | 83 | syllables: 423043 | syllables: 0.042 | 166676.8 | 83823 | [104, 105] |
| European starling | 7 | syllables: 164230 | syllables: 0.577 | 194529.9 | 3805 | [102] |
| Gervais's beaked whale | Unk. | clicks: 1936 | clicks: 0.001 | 2.0 | 1936 | [51, 107] |
| Giant otter | Unk. | syllables: 452 | syllables: 0.68 | 390.4 | 452 | [97] |
| Gibbon | Unk. | syllables: 10333 | syllables: 2.96 | 230400.0 | 128 | [103] |
| Great blue heron | Unk. | syllables: 246 | syllables: 0.138 | 44.1 | 246 | [50, 108] |
| House finch | Unk. | syllables: 248 | syllables: 0.093 | 25.9 | 248 | [50, 108] |
| Human (English) | 40 | words: 283721 | words: 0.205 | 135708.4 | 254 | [94] |
|  |  | phones: 837896 | phones: 0.069 |  |  |  |
| Humpback whale | Unk. | syllables: 2006 | syllables: 1.65 | 6730.8 | 13 | [101] |
| Indigo bunting | Unk. | syllables: 251 | syllables: 0.135 | 36.0 | 251 | [50, 108] |
| Macaque | 8 | coos: 7284 | coos: 0.324 | 2550.9 | 7284 | [23, 106] |
| Marsh wren | Unk. | syllables: 248 | syllables: 0.09 | 23.8 | 248 | [50, 108] |
| Mouse | 4 | syllables: 34124 | syllables: 0.018 | 25277.0 | 133 | [65] |
| Song sparrow | Unk. | syllables: 258 | syllables: 0.105 | 32.8 | 258 | [50, 108] |
| Swamp sparrow | 616 | elements: 97513 | elements: 0.021 | 4571.1 | 1867 | [18, 95] |
| White-rumped munia | 44 | syllables: 109851 | syllables: 0.05 | 17118.5 | 169 | [4] |
| Yellow warbler | Unk. | syllables: 246 | syllables: 0.078 | 21.4 | 246 | [50, 108] |
| Zebra finch | 6 | motifs: 18028 | motifs: 0.443 | 8799.9 | 18028 | [98] |
|  |  | syllables: 65892 | syllables: 0.105 |  |  |  |
| Zebra finch | 46 | elements: 3347 | elements: 0.153 | 1365.0 | 3347 | [99, 100] |

#### 3.1.4 Abstracting sequential organization

Animal vocalizations are not always comprised of single, discrete, temporally-isolated elements (e.g. notes, syllables, or phrases), but often occur as temporally patterned sequences of these elements. The latent projection methods described above can be used to abstract corpora of song elements that can then be used for syntactic analyses [3].

As an example of this, we derived a corpus of symbolically segmented vocalizations from a dataset of Bengalese finch song using latent projections and clustering (Fig 6). Bengalese finch song bouts comprise a small number (~5-15) of highly stereotyped syllables produced in well-defined temporal sequences a few dozen syllables long [4]. We first projected syllables from a single Bengalese finch into UMAP latent space, then visualized transitions between vocal elements in latent space as line segments between points (Fig 6B), revealing highly regular patterns. To abstract this organization to a grammatical model, we clustered latent projections into discrete categories using HDBSCAN. Each bout is then treated as a sequence of symbolically labeled syllables (e.g. *B* → *B* → *C* → *A*; Fig 6D) and the entire dataset rendered as a corpus of transcribed song (Fig 6E). Using the transcribed corpus, one can abstract statistical and grammatical models of song, such as the Markov model shown in Fig 6C or the information-theoretic analysis in Sainburg et al., [3].

##### Sequential organization is tied to transcription method

As we previously noted, hand labels and latent cluster labels of birdsong syllables generally overlap (e.g. Fig 5), but may disagree for a sizable minority of syllables (Table 1). In mice, different algorithmic methods for abstracting and transcribing mouse vocal units (USVs) can result in significant differences between syntactic descriptions of sequential organization [39]. We were interested in the differences between the abstracted sequential organization of birdsong when syllables were labeled by hand versus clustered in latent space. Because we have Bengalese finch datasets that are hand transcribed from two different research groups [8, 57], these datasets are ideal for comparing the sequential structure of algorithmic versus hand-transcribed song.

To contrast the two labeling methods, we first took the two Bengalese finch song datasets, projected syllables into latent space, and visualized them using the hand transcriptions provided by the datasets (Fig 7A,H). We then took the syllable projections and clustered them using HDBSCAN. In both datasets, we find that many individual hand-transcribed syllable categories are comprised of multiple HDBSCAN-labelled clusters in latent space (Fig 7A,B,H,I). To compare the different sequential abstractions of the algorithmically transcribed labels and the hand transcribed labels, we visualized the transitions between syllables in latent space (Fig 7C,J). These visualizations reveal that different algorithmically-transcribed clusters belonging to the same hand-transcribed label often transition to and from separate clusters in latent space. We visualize this effect more explicitly in Fig 7D and K, showing the first-order (incoming and outgoing) transitions between one hand-labeled syllable category (from Fig 7A and H), colored by the multiple HDBSCAN clusters that it comprises (from Fig 7B and I). Thus, different HDBSCAN labels that belong to the same hand-labeled category can play a different role in song-syntax, having different incoming and outgoing transitions. In Fig 7E,F,L,M, this complexity plays out in an abstracted Markov model, where the HDBSCAN-derived model reflects the latent transitions observed in Fig 7D,J more explicitly than the model abstracted from hand-labeled syllables. To further understand why these clusters are labeled as the same category by hand but different categories using HDBSCAN clustering, we show example syllables from each cluster Fig 7G,N. Although syllables from different HDBSCAN clusters look very similar, they are differentiated by subtle yet systematic variation. Conversely, different subsets of the same experimenter-labeled category can play different syntactic roles in song sequences. The syntactic organization in Bengalese finch song is often described using partially observable or hidden Markov models, where the same syllable category plays different syntactic roles dependent on its current position in song syntax [4]. In so far as the sequential organization abstracted from hand labels obscures some of the sequential structure captured by algorithmic transcriptions, our results suggest that these different syntactic roles may be explained by the presence of different syllable categories.

### 3.2 Temporally continuous latent trajectories

Not all vocal repertoires are made up of elements that fall into highly discrete clusters in latent space (Fig 1). For several of our datasets, categorically discrete elements are not readily apparent, making analyses such as those performed in Figure 6 more difficult. In addition, many vocalizations are difficult to segment temporally, and determining what features to use for segmentation requires careful consideration [1]. In many bird songs, for example, clear pauses exist between song elements that enable one to distinguish syllables. In other vocalizations, however, experimenters must rely on less well-defined physical features for segmentation [1, 12], which may in turn invoke a range of biases and unwarranted assumptions. At the same time, much of the research on animal vocal production, perception, and sequential organization relies on identifying “units” of a vocal repertoire [1]. To better understand the effects of temporal discretization and categorical segmentation in our analyses, we considered vocalizations as continuous trajectories in latent space and compared the resulting representations to those that treat vocal segments as single points (as in the previous finch example in Fig. 6). We explored four datasets, ranging from highly discrete clusters of vocal elements (Bengalese finch, Fig. 8), to relatively discrete clustering (European starlings, Fig. 9) to low clusterability (Mouse USV, Fig. 10; Human speech, Fig. 11). In each dataset, we find that continuous latent trajectories capture short and long timescale structure in vocal sequences without requiring vocal elements to be segmented or labeled.

**Figure 8:**
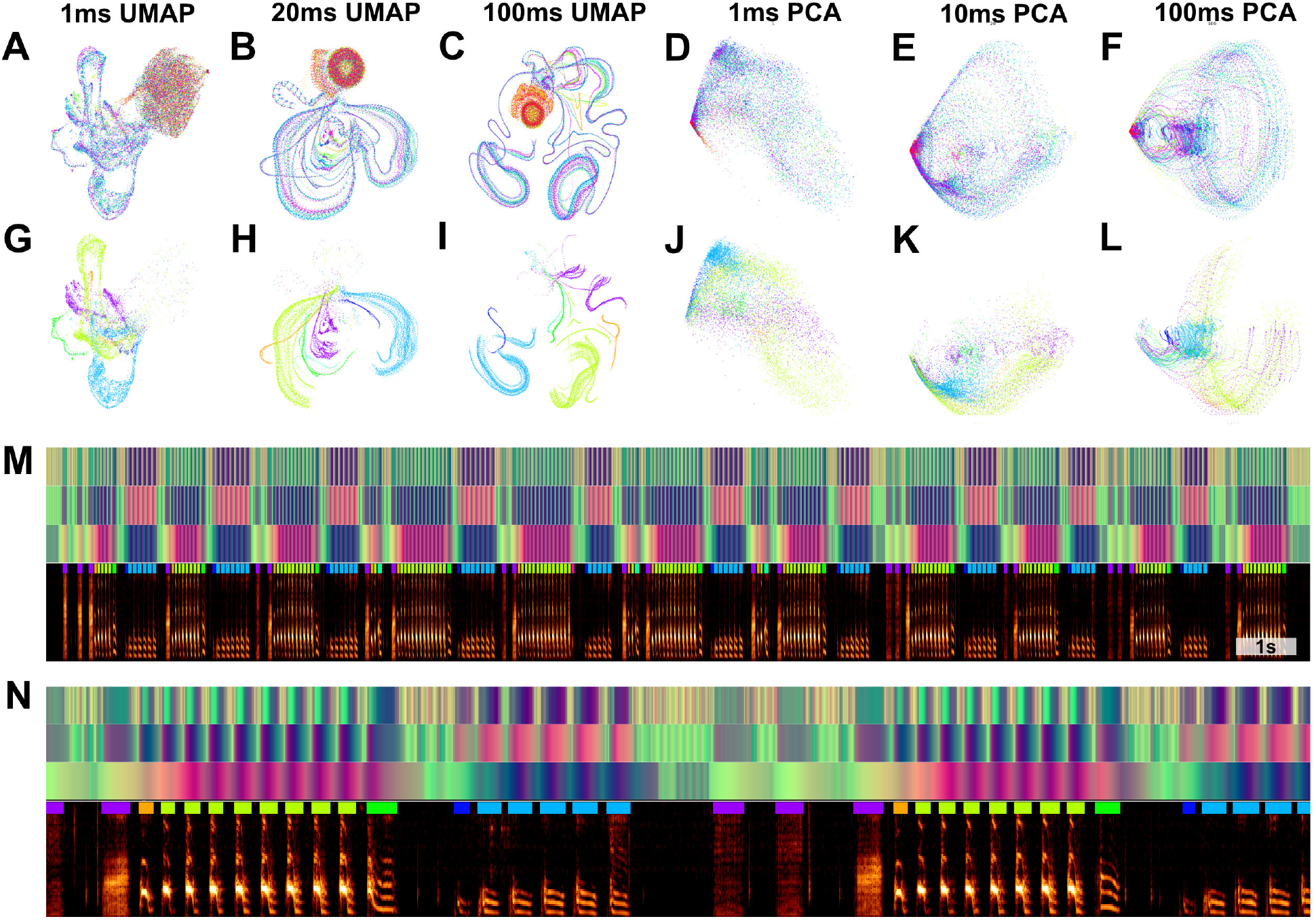
Continuous UMAP projections of Bengalese finch song from a single bout produced by one individual. (A-C) Bengalese finch song is segmented into either 1ms (A), 20ms (B), or 100ms (C) rolling windows of song, which are projected into UMAP. Color represents time within the bout of song. (D-F) The same plots as in (A), projected into PCA instead of UMAP. (G-I) The same plots as (A-C) colored by hand-labeled element categories. (J-L) The same plot as (D-E) colored by hand-labeled syllable categories. (M) UMAP projections projected into colorspace over bout spectrogram. The top three rows are the UMAP projections from (A-C) projected into RGB colorspace to show the position within UMAP space over time as over the underlying spectrogram data. (N) a subset of the bout shown in (M).

**Figure 9:**
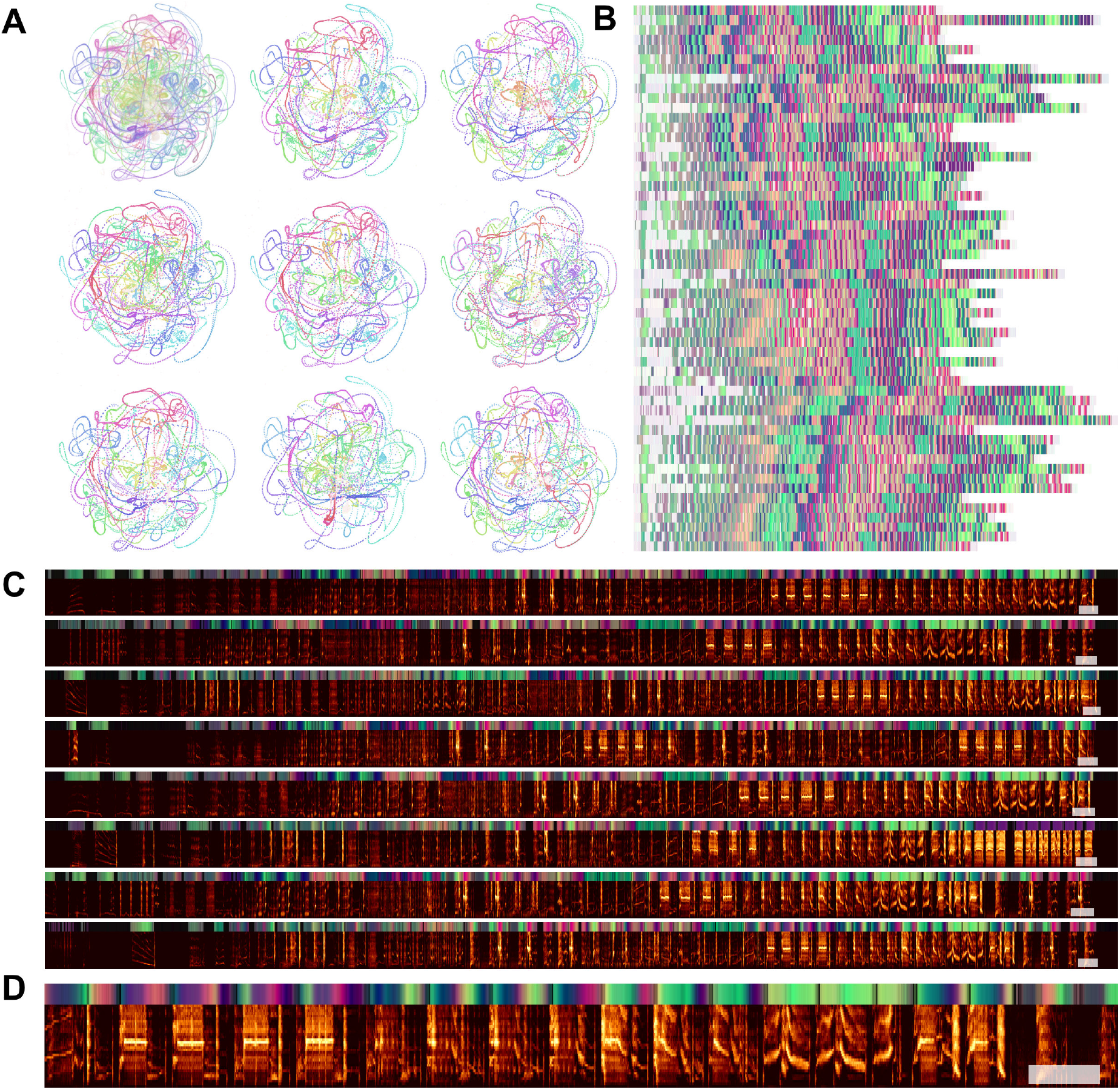
Starling bouts projected into continuous UMAP space. (A) The top left plot is each of 56 bouts of starling song projected into UMAP with a rolling window length of 200ms, color represents time within the bout. Each of the other 8 plots is a single bout, demonstrating the high similarity across bouts. (B) Latent UMAP projections of the 56 bouts of song projected into colorspace in the same manner as Fig 8M. Although the exact structure of a bout of song is variable from rendition to rendition, similar elements tend to occur at similar regions of song and the overall structure is preserved. (C) The eight example bouts from A UMAP colorspace projections above. The white box at the end of each plot corresponds is one second. (D) A zoomed-in section of the first spectrogram in C.

**Figure 10:**
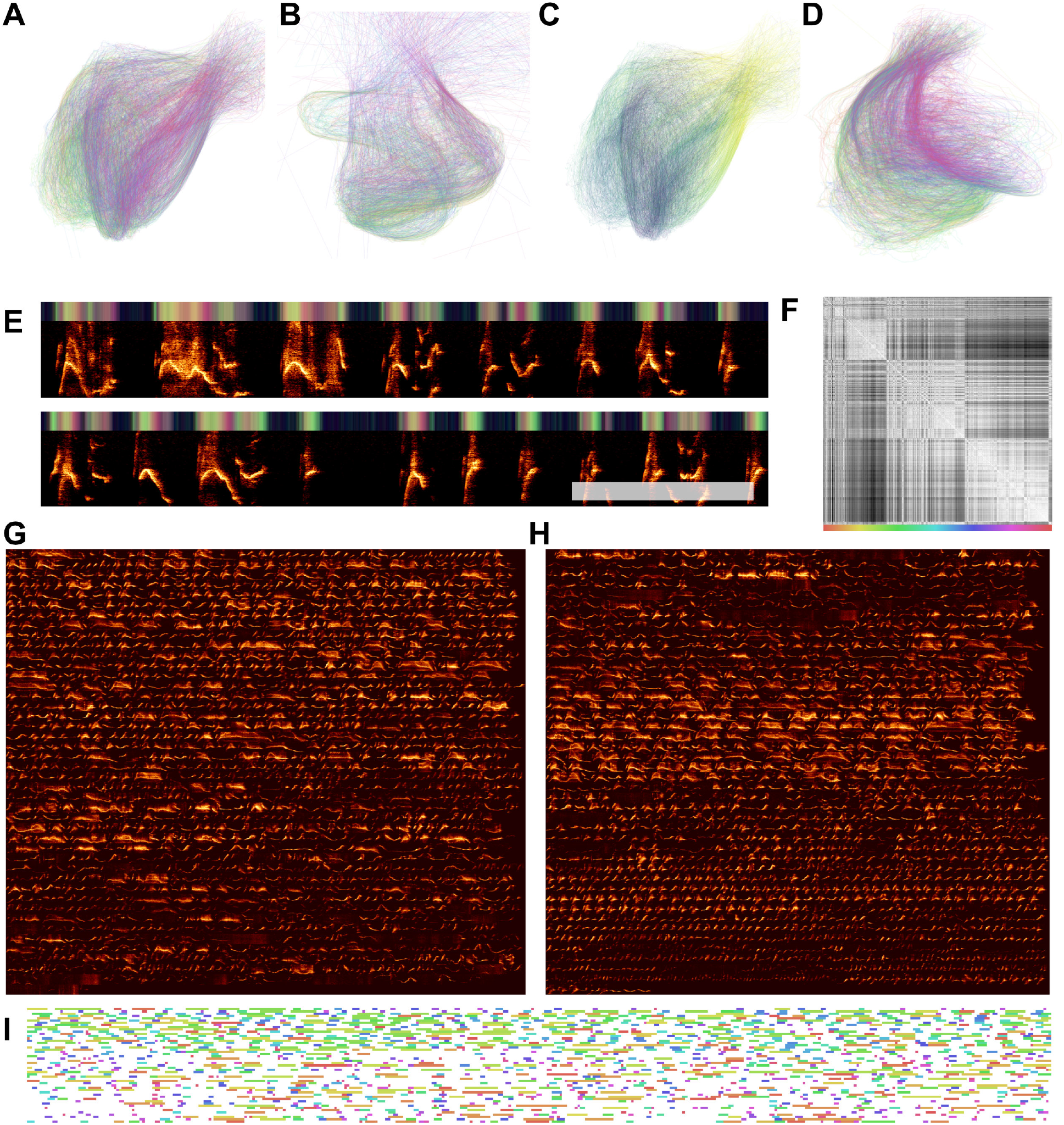
USV patterns revealed through latent projections of a single mouse vocal sequence. (A) Each USV is plotted as a line and colored by its position within the sequence. Projections are sampled from a 5ms rolling window. (B) Projections from a different recording from a second individual using the same method as in (A). (C) The same plot as in A, where color represents time within a USV. (D) The same plot as in (A) but with a 20ms rolling window. (E) An example section of the USVs from (A), where the bar on the top of the plot shows the UMAP projections in colorspace (the first and second USV dimensions are plotted as color dimensions). (F) A similarity matrix between each of 1,590 USVs produced in the sequence visualized in (A), reordered so that similar USVs are closer to one another. (G) Each of the 1,590 USVs produced in the sequence from (A), in order (left to right, top to bottom). (H) The same USVs as in (G), reordered based upon the similarity matrix in (F). (I) The entire sequence from (A) where USVs are color-coded based upon their position in the similarity matrix in (F).

**Figure 11:**
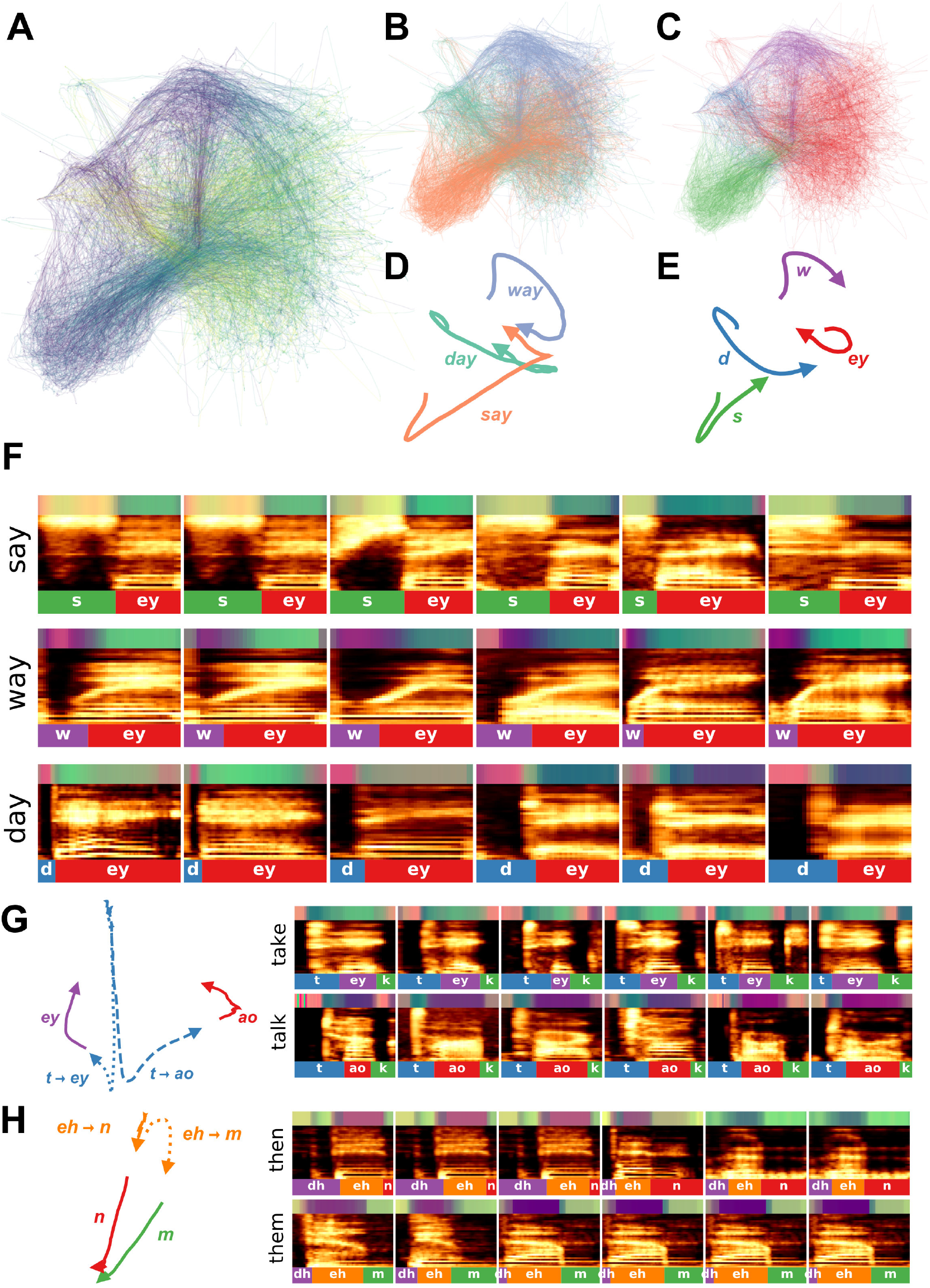
Speech trajectories showing coarticulation in minimal pairs. (A) Utterances of the words ‘day’, ‘say’, and ‘way’ are projected into a continuous UMAP latent space with a window size of 4ms. Color represents time, where darker is earlier in the word. (B) The same projections as in (A) but color-coded by the corresponding word. (C) The same projections are colored by the corresponding phonemes. (D) The average latent trajectory for each word. (E) The average trajectory for each phoneme. (F) Example spectrograms of words, with latent trajectories above spectrograms and phoneme labels below spectrograms. (G) Average trajectories and corresponding spectrograms for the words ‘take’ and ‘talk’ showing the different trajectories for ‘t’ in each word. (H) Average trajectories and the corresponding spectrograms for the words ‘then’ and ‘them’ showing the different trajectories for ‘eh’ in each word.

#### 3.2.1 Comparing discrete and continuous representations of song in the Bengalese finch

Bengalese finch song provides a relatively easy visual comparison between the discrete and continuous treatments of song, because it consists of a small number of unique highly stereotyped syllables (Fig. 8). With a single bout of Bengalese finch song, which contains several dozen syllables, we generated a latent trajectory of song as UMAP projections of temporally-rolling windows of the bout spectrogram (See Projections section). To explore this latent space, we varied the window length between 1 and 100ms (Fig. 8A-L). At each window size, we compared UMAP projections (Fig. 8A-C) to PCA projections (Fig. 8D-F). In both PCA and UMAP, trajectories are more clearly visible as window size increases across the range tested, and overall the UMAP trajectories show more well-defined structure than the PCA trajectories. To compare continuous projections to discrete syllables, we re-colored the continuous trajectories by the discrete syllable labels obtained from the dataset. Again, as the window size increases, each syllable converges to a more distinct trajectory in UMAP space (Fig. 8G-I). To visualize the discrete syllable labels and the continuous latent projections in relation to song, we converted the 2D projections into colorspace and show them as a continuous trajectory alongside the song spectrograms and discrete labels in Figure 8M,N.

#### 3.2.2 Latent trajectories of European starling song

European starling song provides an interesting case study for exploring the sequential organization of song using continuous latent projections because starling song is more sequentially complex than Bengalese finch song, but is still highly stereotyped and has well-characterized temporal structure. European starling song is comprised of a large number of individual song elements, usually transcribed as ‘motifs’, that are produced within a bout of singing. Song bouts last several tens of seconds and contain many unique motifs grouped into three broad classes: introductory whistles, variable motifs, and high-frequency terminal motifs [61]. Motifs are variable within classes, and variability is affected by the presence of potential mates and seasonality [62, 63]. Although sequentially ordered motifs are usually segmentable by gaps of silence occurring when starlings are taking breaths, segmenting motifs using silence alone can be difficult because pauses are often short and bleed into surrounding syllables [64]. When syllables are temporally discretized, they are relatively clusterable (Fig 1), however syllables tend to vary somewhat continuously (Fig 9D). To analyze starling song independent of assumptions about segment (motif) boundaries and element categories, we projected bouts of song from a single male European starling into UMAP trajectories using the same methods as in Figure 8.

We find that the broad structure of song bouts are highly repetitive across renditions, but contain elements within each bout that are variable across bout renditions. For example, in Figure 9A, the top left plot is an overlay showing the trajectories of 56 bouts performed by a single bird, with color representing time within each bout. The eight plots surrounding it are single bout renditions. Different song elements are well time-locked as indicated by a strong hue present in the same regions of each plot. Additionally, most parts of the song occur in each rendition. However, certain song elements are produced or repeated in some renditions but not others. To illustrate this better, in Fig 9B, we show the same 56 bouts projected into colorspace in the same manner as Fig 8M,N, where each row is one bout rendition. We observe that, while each rendition contains most of the same patterns at relatively similar times, some patterns occur more variably. In Fig 9C and D we show example spectrograms corresponding to latent projections in Fig 9A, showing how the latent projections map onto spectrograms.

Quantifying and visualizing the sequential structure of song using continuous trajectories rather than discrete element labels is robust to errors and biases in segmenting and categorizing syllables of song. Our results show the potential utility of continuous latent trajectories as a viable alternative to discrete methods for analyzing song structure even with highly complex, many-element, song.

#### 3.2.3 Latent trajectories and clusterability of mouse USVs

House mice produce ultrasonic vocalizations (USVs) comprising temporally discrete syllable-like elements that are hierarchically organized and produced over long timescales, generally lasting seconds to minutes [65]. When analyzed for temporal structure, mouse vocalizations are typically segmented into temporally-discrete USVs and then categorized into discrete clusters [1, 39, 65–67] in a manner similar to syllables of birdsong. As Figure 1 shows, however, USVs do not cluster into discrete distributions in the same manner as birdsong. Choosing different arbitrary clustering heuristics will therefore have profound impacts on downstream analyses of sequential organization [39].

We sought to better understand the continuous variation present in mouse USVs, and explore the sequential organization of mouse vocalizations without having to categories USVs. To do this, we represented mouse USVs as continuous trajectories (Fig 10E) in UMAP latent space using similar methods as with starlings (Fig. 8) and finches (Fig. 9). In Figure 10, we use a single recording of one individual producing 1,590 (Fig. 10G) USVs over 205 seconds as a case study to examine the categorical and sequential organization of USVs. We projected every USV produced in that sequence as a trajectory in UMAP latent space (Fig. 10A,C,D). Similar to our observations in Figure 1I using discrete segments, we do not observe clear element categories within continuous trajectories, as observed for Bengalese finch song (e.g. Fig 8I).

To explore the categorical structure of USVs further, we reordered all of the USVs in Figure 10G by the similarity of their latent trajectories (Fig. 10F) and plotted them side-by-side (Fig. 10H). Both the similarity matrix of the latent trajectories (Fig. 10F) and the similarity-reordered spectrograms (Fig. 10H) show that while some USVs are similar to their neighbors, no highly stereotyped USV categories are observable.

Although USVs do not aggregate into clearly discernible, discrete clusters, the temporal organization of USVs within the vocal sequence is not random. Some latent trajectories are more frequent at different parts of the vocalization. In Figure 10A, we color-coded USV trajectories according to each USV’s position within the sequence. The local similarities in coloring (e.g., the purple and green hues) indicate that specific USV trajectories tend to occur in distinct parts of the sequence. Arranging all of the USVs in order (Fig. 10G) makes this organization more evident, where one can see that shorter and lower amplitude USVs tend to occur more frequently at the end of the sequence. To visualize the vocalizations as a sequence of discrete elements, we plotted the entire sequence of USVs (Fig. 10I), with colored labels representing the USV’s position in the reordered similarity matrix (in a similar manner as the discrete category labels in Fig. 6E. In this visualization, one can see that different colors dominate different parts of the sequence, again reflecting that shorter and quieter USVs tend to occur at the end of the sequence.

#### 3.2.4 Latent trajectories of human speech

Discrete elements of human speech (i.e. phonemes) are not spoken in isolation, and their acoustics are influenced by neighboring sounds, a process termed co-articulation. For example, when producing the words ‘day’, ‘say’, or ‘way’, the position of the tongue, lips, and teeth differ dramatically at the beginning of the phoneme ‘ey’ due to the preceding ‘d’, ‘s’, or ‘w’ phonemes, respectively. This results in differences in the pronunciation of ‘ey’ across words (Fig 11F). Co-articulation explains much of the acoustic variation observed within phonetic categories. Abstracting to phonetic categories therefore discounts much of this context-dependent acoustic variance.

We explored co-articulation in speech, by projecting sets of words differing by a single phoneme (i.e. minimal pairs) into continuous latent spaces, then extracted trajectories of words and phonemes that capture sub-phonetic context-dependency (Fig. 11). We obtained the words from the same Buckeye corpus of conversational English used in Figures 1, 4, and 18. We computed spectrograms over all examples of each target word, then projected sliding 4-ms windows from each spectrogram into UMAP latent space to yield a continuous vocal trajectory over each word (Fig. 11). We visualized trajectories by their corresponding word and phoneme labels (Fig. 11B,C) and computed the average latent trajectory for each word and phoneme (Fig. 11D,E). The average trajectories reveal context-dependent variation within phonemes caused by coarticulation. For example, the words ‘way’, ‘day’, and ‘say’ each end in the same phoneme (‘ey’; Fig. 11A-F), which appears as an overlapping region in the latent space (the red region in Fig 11C). The endings of each average word trajectory vary, however, indicating that the production of ‘ey’ differs based on its specific context (Fig 11D). The difference between the production of ‘ey’ can be observed in the average latent trajectory over each word, where the trajectories for ‘day’ and ‘say’ end in a sharp transition, while the trajectory for ‘way’ is more smooth (Fig 11D). These differences are apparent in figure 11F which shows examples of each word’s spectrogram accompanied by its corresponding phoneme labels and color-coded latent trajectory. In the production of ‘say’ and ‘day’ a more abrupt transition occurs in latent space between ‘s’/‘d’ and ‘ey’, as indicated by the yellow to blue-green transitions above spectrograms in ‘say’ and the pink to blue-green transition above ‘day’. For ‘way’, in contrast, a smoother transition occurs from the purple region of latent space corresponding to ‘w’ to the blue-green region of latent space corresponding to ‘ey’.

Latent space trajectories can reveal other co-articulations as well. In Figure 11G, we show the different trajectories characterizing the phoneme ‘t’ in the context of the word ‘take’ versus ‘talk’. In this case, the ‘t’ phoneme follows a similar trajectory for both words until it nears the next phoneme (‘ey’ vs. ‘ao’), at which point the production of ‘t’ diverges for the different words. A similar example can be seen for co-articulation of the phoneme ‘eh’ in the words ‘them’ versus ‘then’ (Fig. 11H). These examples show the utility of latent trajectories in describing sub-phonemic variation in speech signals in a continuous manner rather than as discrete units.

### 3.3 Probing the latent space with neural networks and generative models

All of the examples shown here so far use latent models to capture variation in complex vocal signals. These methods enable new forms of visualization, help improve understanding of the structure of animal communication, and yield an unbiased (or at least less biased) feature set from which analyses of vocal communication can be performed. An even more powerful application of latent models, however, is in generating novel stimuli [30, 31, 68]. Using generative latent models of animal vocalizations, one can systematically probe perceptual and neural representations of vocal repertoires in complex natural feature spaces. To do this, latent models must be bidirectional: in addition to projecting vocalizations into latent space, they must also sample from latent space to generate novel vocalizations. That is, where dimensionality reduction only needs to project from vocalization space (*X*) to latent space (*Z*), *X*→*Z*, generativity requires bidirectionality: *X*↔*Z*. In the following section we discuss and explore the relative merits of a series of neural network architectures that are designed to both reduce dimensionality and generate novel data.

#### 3.3.1 Neural network architectures

While much of the attention paid to deep neural networks in the past decade had focused on advances in applications of supervised learning such as image classification or speech recognition [13], neural networks have also made substantial advancements in the fields of dimensionality reduction and data representation [14, 37]. Like UMAP, deep neural networks can be trained to learn reduced-dimensional, compressive, representations of complex data using successive layers of nonlinearity. They do so using network architectures and error functions that encourage the compressive representation of complex data. Here we survey a set of network architectures and show their applicability to modeling animal vocalizations.

##### Autoencoders

Perhaps the simplest example of a neural network that can both reduce dimensionality (*X* → *Z*) and generate novel data (*Z* → *X*) is the autoencoder (AE; [34]). AEs comprise two subnetworks, an *encoder* which translates from *X* → *Z* and a *decoder* which translates from *Z* → *X* (Fig. 12A). The network is trained on a single error function: to reconstruct in *X* as well as possible. Because this reconstruction passes through a reduced-dimensional latent layer (*Z*), the encoder learns an encoding in *Z* that compressively represents the data, and the decoder learns to generate data back into *X* from compressed projections in *Z*. Both sub-networks contain stacked layers of non-linear artificial neurons that learn either more compressive or more generalizable representations of their inputs (*X* or *Z*, respectively) as depth in the sub-network increases [69, 70].

**Figure 12:**
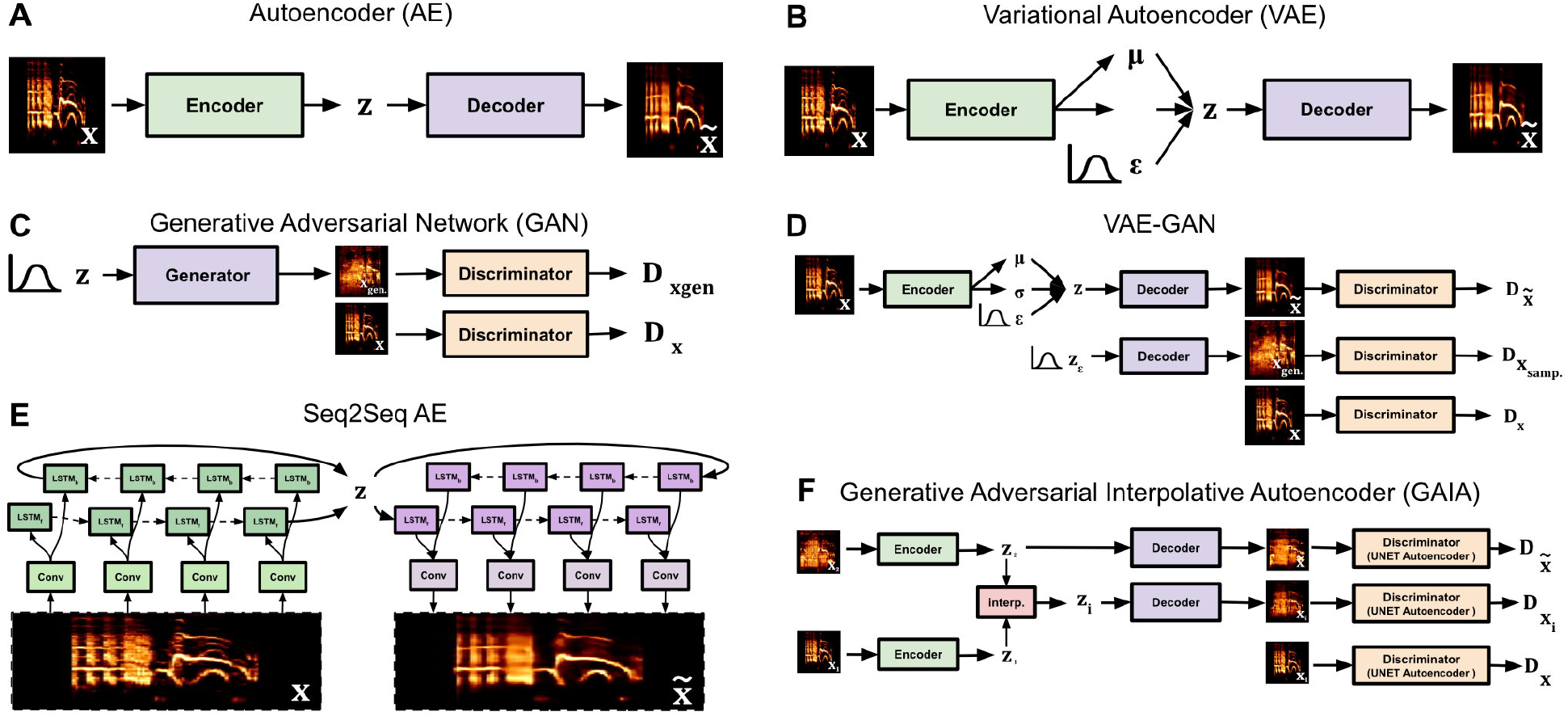
Neural network architectures, latent projections, and reconstructions. (A) Autoencoder. (B) Variational autoencoder. (C) Generative adversarial network. (D) Variational Autoencoder / Generative Adversarial Network (VAE-GAN) (E) Seq2Seq autoencoder. (F) Generative Adversarial Interpolative autoencoder. Note that the Seq2Seq autoencoder follows the same general architecture as A, with the encoder and decoder shown in more detail. The Multidimensional Scaling Autoencoder (MD-AE, see text) also uses the same general architecture as (A) with an additional loss function from a base autoencoder.

##### Generative Adversarial Networks

A second, more recent network architecture is the Generative Adversarial Network (GAN; [37]). GANs are so-named In the because they are composed of two networks, the generator, and the discriminator, which act adversarially against one another to generate data in *X* (Fig. 12C). The generator acts similarly to the decoder in an AE, projecting data from *Z* into *X*, however, instead of reconstructing data, the generator samples from a pre-defined distribution (e.g. uniform or Gaussian) in *Z* and attempts to construct a sample in *X*, with the goal of fooling the discriminator into classifying the generated data as being real data. The discriminator meanwhile, is given both real data and data generated by the generator and is tasked with differentiating between the two. As both networks learn, the discriminator gets better at differentiating between real and fake, and the generator gets better at producing fakes that are less distinguishable from real data. Notably, GANs are unidirectional in that they can only translate from *Z* → *X*, and not *X* → *Z*, meaning that they generate data but do not perform dimensionality reduction.

##### Generative models and Variational Autoencoders

GANs belong to a more specific class of models, *generative models*, in which the joint probability of the latent distribution and the data distribution are modeled directly [71]. AEs are not generative models by default, but the Variational Autoencoder (VAE), an autoencoder with an additional regularization loss to encode data into a predefined (usually Gaussian) latent distribution [72], is a generative model (Fig. 12B). Generative models are often preferable because they can be sampled from probabilistically in latent space.

##### VAE-GANs

AEs, VAEs, and GANs each possess attributes that are a combination of beneficial or detrimental for modeling animal vocalizations. A primary detriment of GANs is that they are not bidirectional, i.e. they do not translate from *X* → *Z*. A primary detriment of AEs and VAEs is that they are trained directly on reconstructions in *X*, resulting in reconstructions of compressed syllables that tend to look blurry and smoothed-over (Fig. 14A,B). Because GANs are not trained on a reconstruction loss but are instead trained to fool a discriminator, generated data better match the sharper contrast of the original data (Fig. 14D). One solution to overcome the blurred reconstructions of AEs and the unidirectionality of GANs is to combine them. One such example is the VAE-GAN [36]. VAE-GANs are comprised of an encoder, decoder, and discriminator (Fig. 12D). The discriminator is trained on a GAN loss to differentiate between sampled and real data. The encoder is trained on both a reconstruction loss (in GAN latent space) and a VAE latent regularization. The decoder is trained both on a reconstruction loss and on the generator GAN loss. The effect of each of these networks and losses in combination is a network that can reconstruct data like a VAE, but where reconstructions are less blurry, like a GAN.

##### Generative Adversarial Interpolative Autoencoders and Multidimensional Scaling Autoencoders

One detriment of the dimensionality reduction component of VAEs and GANs (and VAE-GANs) is that latent projections are forced into a pre-defined latent distribution (e.g. a Gaussian distribution), potentially discarding important structure in the dataset. In contrast, UMAP projections of vocal repertoires are generally non-Gaussian and differ between datasets (Fig. 1), presumably retaining some structure in the data that is otherwise lost with a VAE or GAN. For this reason, we introduce two other network architectures: the multidimensional scaling autoencoder ([35, 68, 73], and the generative adversarial interpolative autoencoder (GAIA; Fig. 12F; [35]). The MDS-AE is an AE that preserves structure in latent representations using an additional multidimensional scaling [35, 74] regularization term so that input relationships in *X* are preserved in *Z*. GAIA is an autoencoder that uses an adversarial loss function to explicitly train data generated from interpolations between projections in *Z* to be indifferentiable from the original data, thus improving interpolations without forcing a predefined latent distribution. Adversarial training on interpolations has previously been shown to produce data representations that have greater utility than AEs of VAEs in some downstream tasks [75].

##### Convolutional and Recurrent layers

Sub-networks such as the encoder and decoder in an AE can be comprised of several different forms of layers and connections. The simplest form is the fully-connected layer, in which each artificial neuron in a given layer is connected to every artificial neuron in the following layer. These architectures are computationally simple but costly, because of the large number of connections in the network. More typically, convolutional layers are used with two-dimensional data like images or spectrograms of syllables. Convolutional layers are loosely motivated by the architecture of primate early sensory cortex [13], and have “receptive fields” that extend over only a subset of the data (e.g. a time-frequency range in a spectrogram). Recurrent layers respect temporal relationships and are more typically used in sequentially organized data, like speech recognition and generation [77, 78] because they contain artificial neurons that learn to preserve temporal relationships (Fig. 12E). Because recurrent networks unfold over time, their latent projections can be treated as trajectories much like the UMAP trajectories in Figs. 8, 9, and 10.

##### Network comparisons

Each network architecture confers its own advantages in representing animal vocalizations. We compared different architectures by projecting the same dataset of canary syllables with equivalent (convolutional) sub-networks and a 2D latent space (Fig. 13; See Neural Networks). As expected, different network architectures produce different latent projections of the same dataset. The VAE and VAE-GAN produce latent distributions that are more Gaussian (Fig. 13E,G) than the MDS-AE and AE (Fig. 13A,C). We then sampled uniformly from latent space to visualize the different latent representations learned by each network (Fig. 13B,D,F,H). Additionally, we trained several network architectures on higher dimensional spectrograms of European starlings syllables with a 128-dimensional latent space (Fig. 14). We plotted reconstructions of syllables as J-diagrams [76] which show both reconstructions and morphs generated through latent interpolations between syllables [30, 31]. Across networks, we observe that syllables generated with AEs (Fig 14A,B) appear more smoothed over, while reconstructions using adversarial-based networks appear less smoothed over but reconstructed syllables match the original syllables less closely (Fig 14C,D).

**Figure 13:**
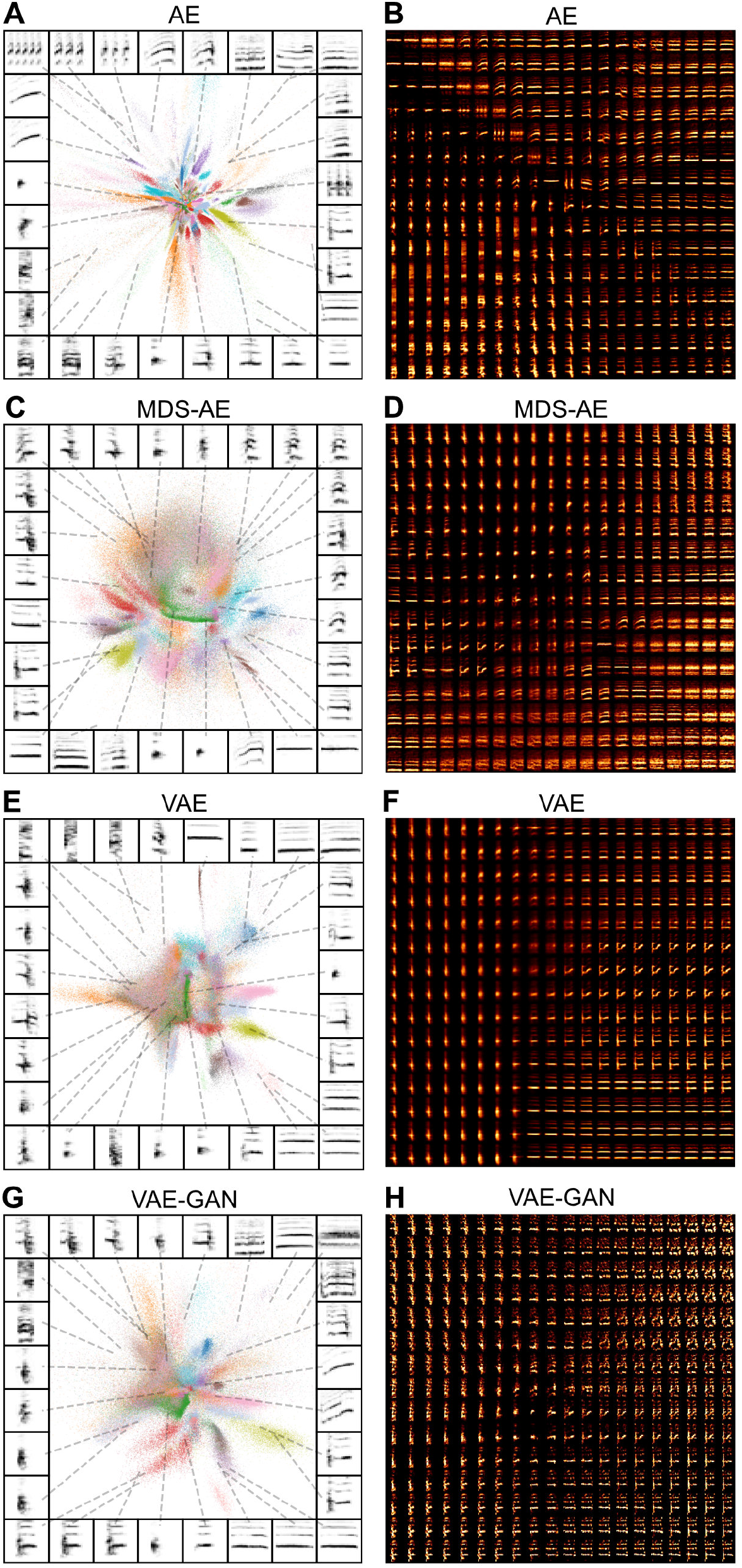
Latent projections and reconstructions of canary syllables.(A-D) Latent projections of syllables, where song phrase category is colored. (E-H) Uniform grids sampled the latent spaces depicted in (A-D).

**Figure 14:**
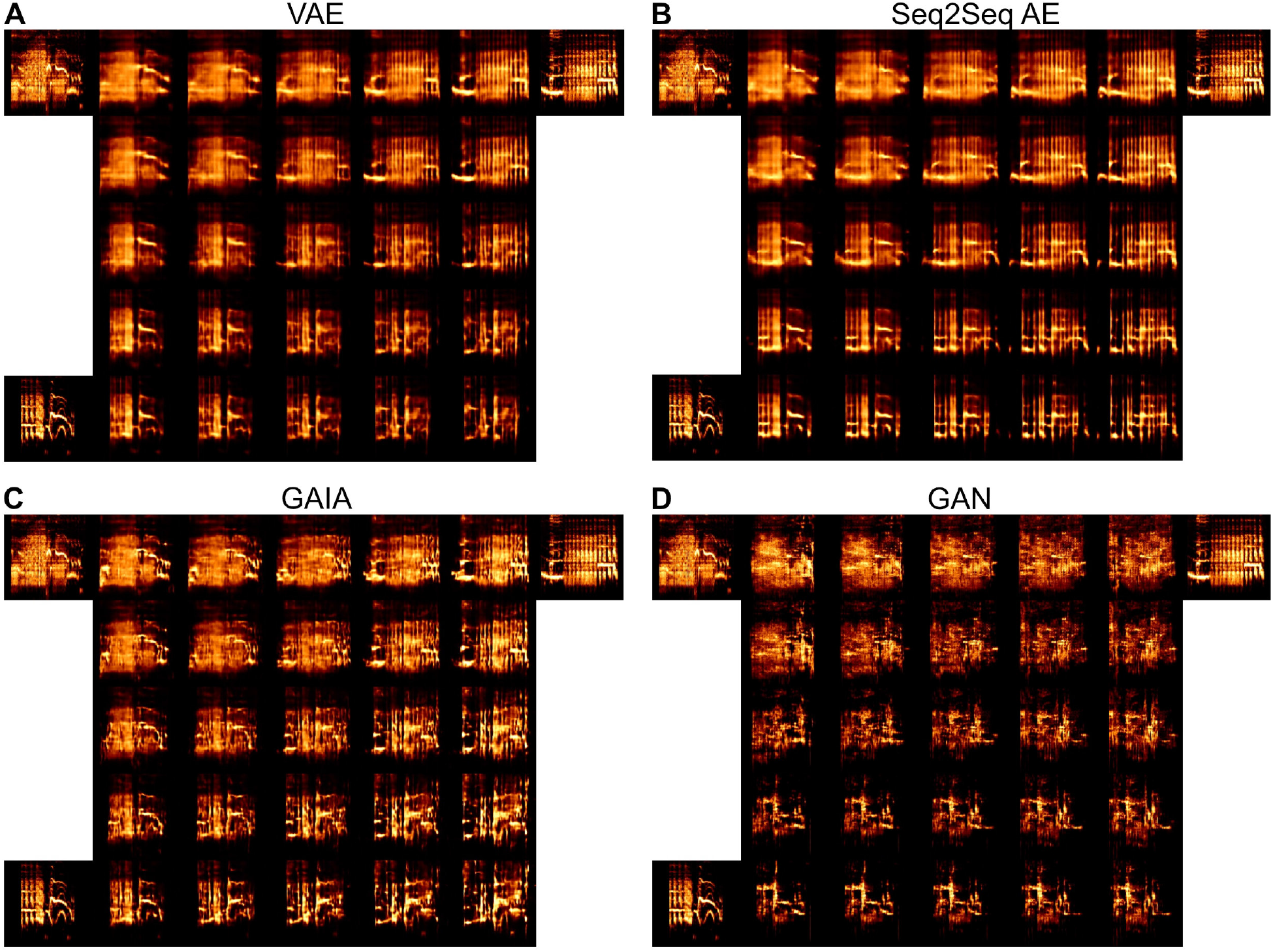
Latent interpolations of European starling syllables. J-Diagrams [76] of syllables reconstructed using different network architectures. The original syllables are shown in the top left (*a*), top right (*b*), and bottom left (*c*) corners, alongside reconstructions. Each other syllable is an interpolation between those syllables, where the bottom right is equal to *b* + *c − a*.

#### 3.3.2 Probing perceptual and neural representations of vocal repertoires

Psychoacoustic studies with animals, such as those common to auditory neuroscience, have focused traditionally on highly simplified stimulus spaces. The utility of simplified stimulus spaces in systems neuroscience is limited however, because many brain regions are selectively or preferentially responsive to naturalistic stimuli, like conspecific or self-generated vocalizations [79, 80]. In contrast, stimuli generated using deep neural networks can be manipulated systematically while still retaining their complex, ecologically-relevant acoustic structure. To demonstrate the utility of generative neural networks for perceptual and neural experiments, we trained European starlings on a two-alternative choice behavioral task in which starlings classified morphs of syllables generated through interpolations in the latent space of a VAE. We then recorded extracellular spiking responses from songbird auditory cortical neurons during passive playback of the morph stimuli. The data presented here is a small subset of the data from a larger ongoing experiment on context-dependency and categorical decision-making [30].

##### Behavioral paradigm

We trained a convolutional VAE on syllables of European starling song and produced linear interpolations between pairs of syllables in the same manner as was shown in Figure 14A. We sampled six acoustically distinct syllables of song, three were arbitrarily assigned to one response class, and three to another. Interpolations between syllables across each category yield nine separate motif continua along which each training motif gradually morphs into one associated with the opposite response. Spectrograms generated from the interpolations were then reconstructed into waveforms using the Griffin-Lim algorithm [81]. This produced nine smoothly varying syllable-morphs (Fig. 15B,E) that we used as playback stimuli in our behavioral experiment.

**Figure 15:**
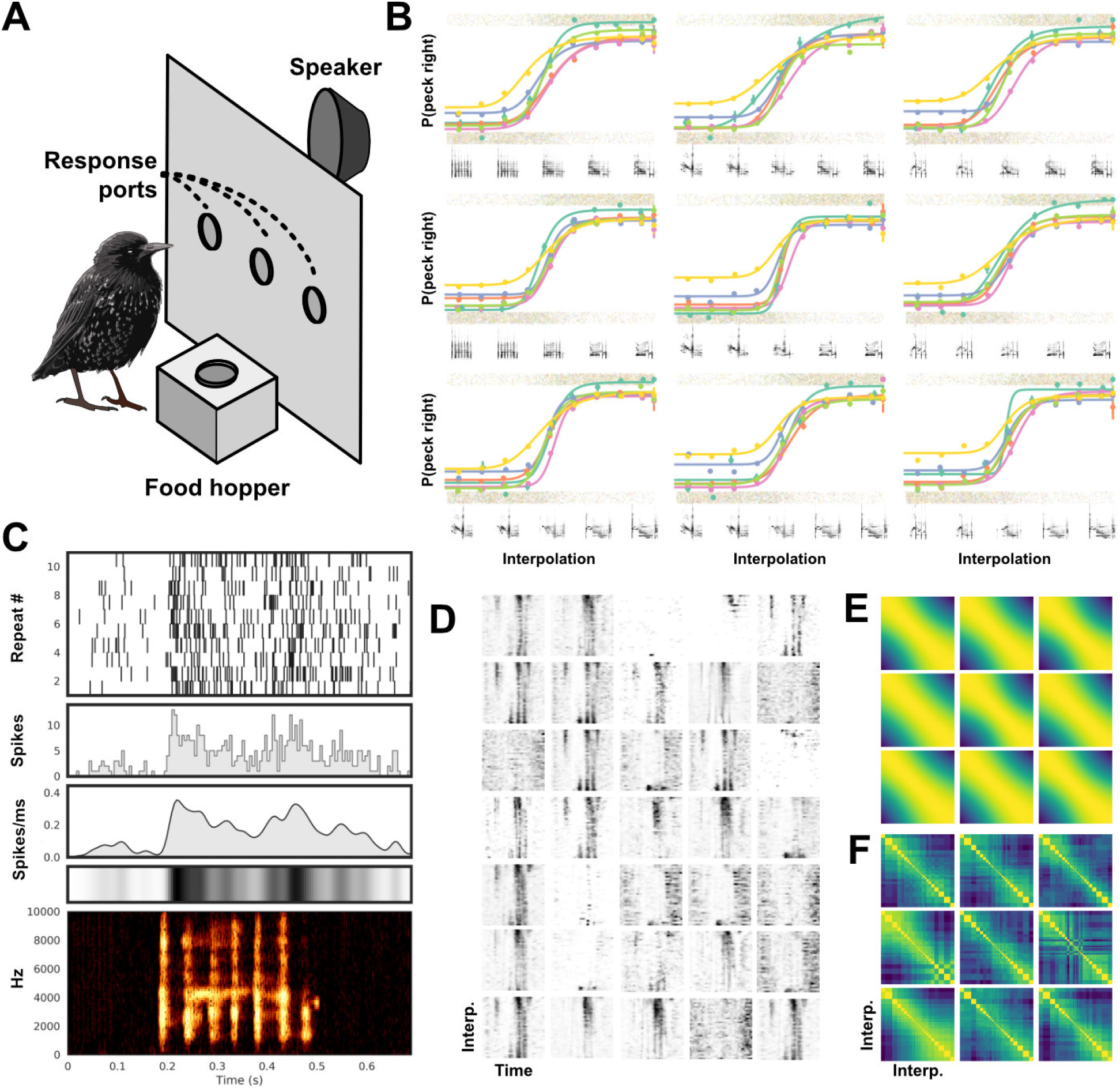
Example behavioral and physiological experiment using VAE generated morphs of European starling song. (A) Birds are trained on a two-alternative choice operant conditioning task where behavioral responses (pecking in response ports) are either rewarded with food or punished with lights-out. (B) Fit psychometric functions for behavioral responses of six birds (differentiated by color) to nine different interpolations (each subplot). The mean and standard error of binned responses (gray points jittered to show distribution) is plotted over top of the fit psychometric functions. Six spectrograms sampled from each morph continuum are shown below the corresponding psychometric function. (C) Responses of a single example neuron recorded from CM in a trained starling in response to one motif sampled from the training set. The plots from top to bottom show a spike raster, the corresponding Peri-Stimulus Time Histogram (PSTH, 5ms bins), and the PSTH convolved with a Gaussian (*σ*=10ms), shown as a lineplot and a colorbar. At the bottom is the spectrogram of the motif. (D) Responses from 35 simultaneously recorded neurons to a single motif continuum. Each plot is a neuron, and each row is a colorbar of the Gaussian convolved PSTH (as in C) to one motif along the continuum. The x-axis is time. (E) Cosine similarity matrix for the spectrogram of each motif along all nine possible morph continua. (F) Cosine similarity matrices of the population responses of all 35 neurons from (D) over each of the nine interpolations as in (B and E). Stimuli near the category boundary are sampled at a higher resolution.

**Figure 16:**
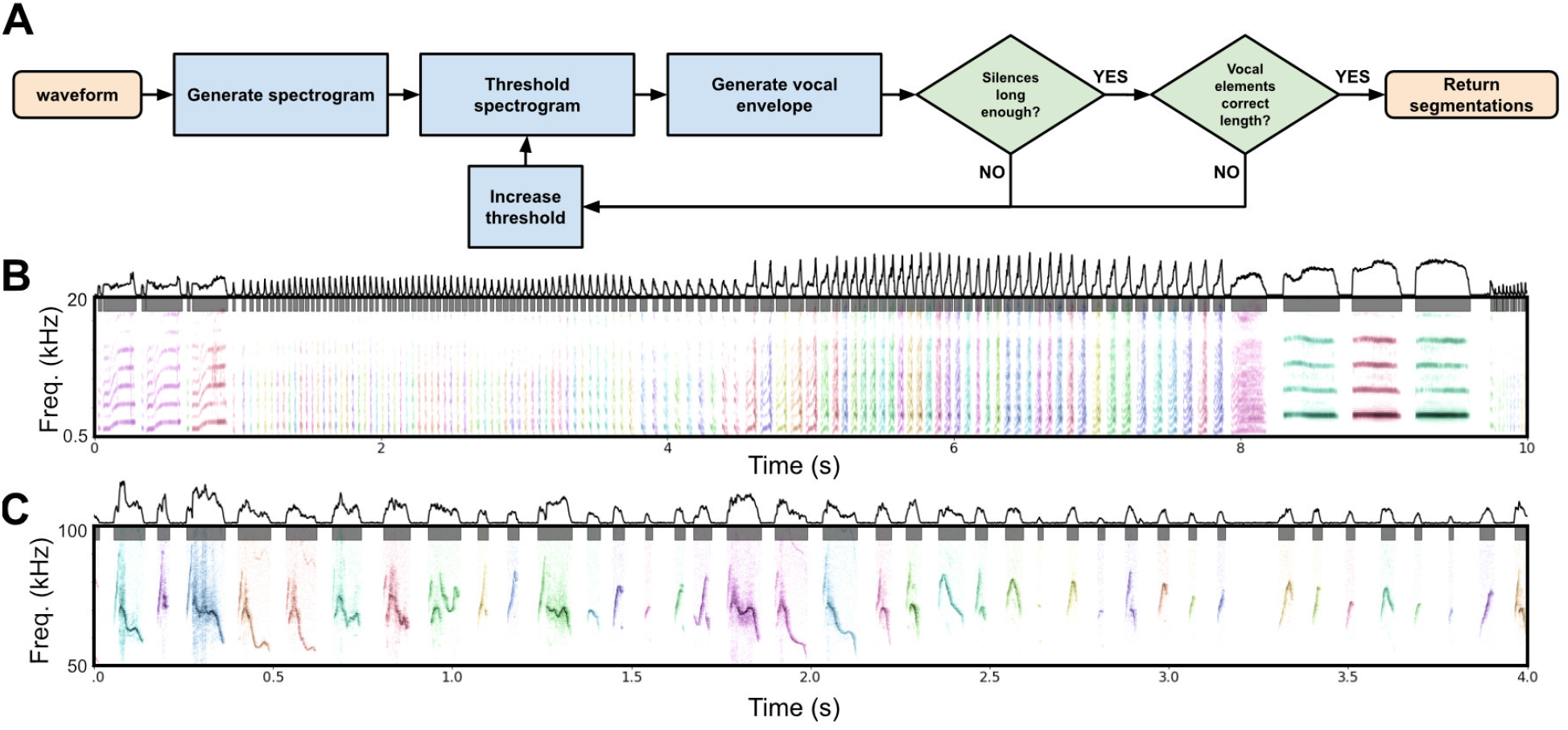
Segmentation algorithms. (A) The dynamic threshold segmentation algorithm. The algorithm dynamically a noise threshold based upon the expected amount of silence in a clip of vocal behavior. Syllables are then returned as continuous vocal behavior separated by noise. (B) The segmentation method from (A) applied to canary syllables. (C) The segmentation method from (A) applied to mouse USVs.

**Figure 17:**
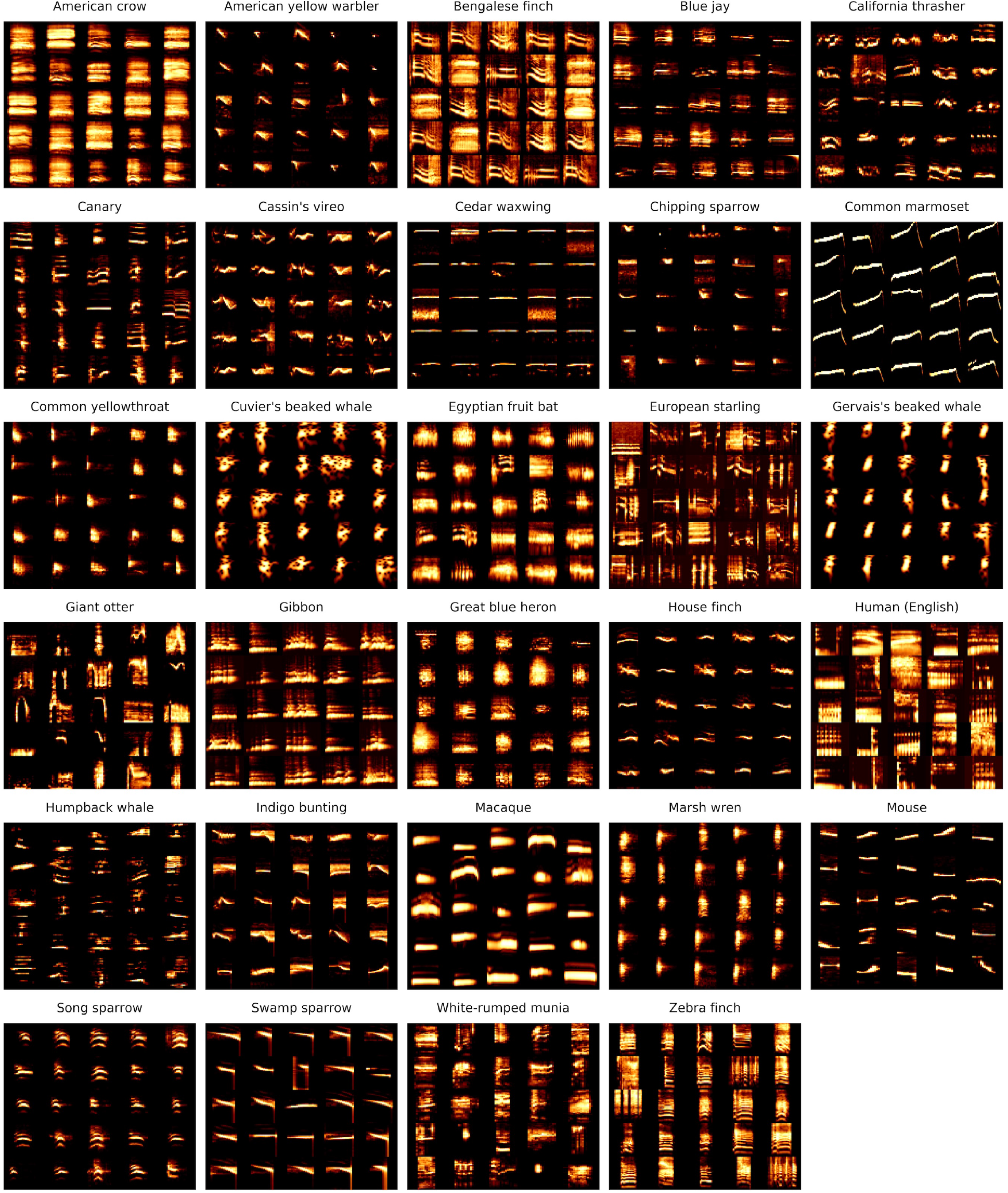
Example vocal elements from each of the species used in this paper.

**Figure 18:**
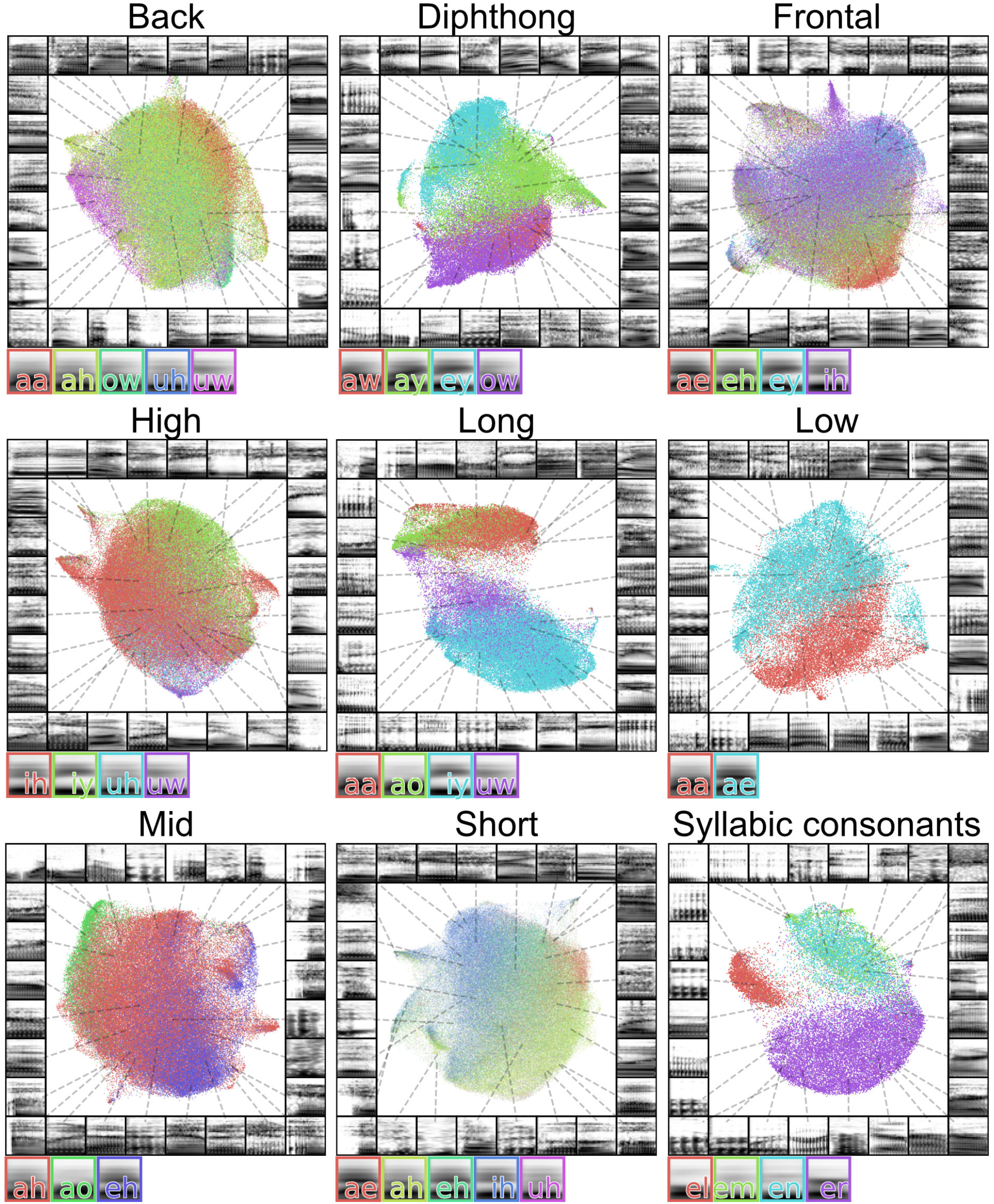
Latent projections of vowels. Each plot shows a different set of vowels grouped by phonetic features. The average spectrogram for each vowel is shown to the right of each plot.

Using an established operant conditioning paradigm ([82]; Fig. 15A), we trained six European starlings to associate each morph with a peck to either the left or right response port to receive food. The midpoint in each motif continuum was set as the categorical boundary. Birds learned to peck the center response port to begin a trial and initiate presentation of a syllable from one of the morph continua, which the bird classifies by pecking into either the left or right response port. Correct classification led to intermittent reinforcement with access to food; incorrect responses triggered a brief (1-5 second) period during which the house light was extinguished and food was inaccessible. Each bird learned the task to a level of proficiency well above chance (~80% - 95%; chance=50%), and a psychometric function of pecking behavior (Fig. 15B) could be extracted, showing that starlings respond to the smooth variation of complex natural stimuli generated from latent interpolations. This mirrors behavioral results on simpler and more traditional parametrically controlled stimuli (e.g. [83, 84]), and provides a proof of concept that ecologically relevant stimuli generated with neural networks can be a viable alternative to simpler stimuli spaces in psychoacoustic research. Because some neurons are selectively responsive to complex or behaviorally relevant stimuli [79, 80], this approach has the potential to open up new avenues for investigating neural representations of auditory objects in ways that more traditional methods cannot.

##### Neural recordings

Songbird auditory forebrain regions such as the caudal medial nidopallium (NCM) and the caudal mesopallium (CM) contain many neurons that respond selectively with a change in spiking activity to conspecific vocalizations, including song [85]. We asked whether VAE-generated artificial stimuli could elicit responses in a song-selective auditory region (CM), and if so whether such responses vary smoothly along morph continua. To do this, we presented the VAE-generated syllable morphs (from the behavioral experiment) to trained, lightly anesthetized birds, while recording from extracellularly on a 32-channel silicon electrode inserted into CM using established methods [86].

An example playback experiment using the same morph stimuli as in the operant conditioning behavior is shown in Figure 15. We then extracted spikes (i.e. putative action potentials) and clustered them into units (i.e. putative neurons) using the MountainSort spike sorting algorithm [87]. For each unit, and in response to each stimulus in each morph, we computed the PSTH of that unit’s response to the stimulus over repeated presentations, then convolved the PSTH with a 5ms Gaussian kernel to get an instantaneous spike rate vector over time for each of the stimuli (Fig. 15C). Figure 15D shows an example of the spike rate vector (as in 15C) for each of the stimuli in a single morph continuum for each of 35 putative simultaneously recorded neurons extracted from one recording site. Figure 15F shows the similarity between neural population responses (as in 15D) for all nine morph continua. We observe that neural responses at both the single neuron and population level (Figs 15D, F) vary smoothly in response to the smoothly varying syllable morphs (Figs 15F). Thus, high-level auditory regions appear to carry a nearly continuous representation of the latent spaces described here, functionally mirroring the responses to simpler features like fundamental frequency observed in many lower-order neurons (e.g. [88]).

## 4 Discussion

We sampled a diverse array of animal vocal communication signals and explored a set of techniques to visualize, systematically analyze, and generate vocalizations through latent representational spaces. Using these techniques we showed that variability exists in the compressed latent space representations of vocal elements across animal species, including songbirds, primates, rodents, bats, and cetaceans (Fig. 1). In general, songbirds tend to produce signals that cluster discretely in latent space, whereas mammalian vocalizations are more uniformly distributed. This observation deserves much closer attention with even more species. We also showed that complex features of datasets, such as individual identity (Fig. 2), species identity (Fig. 3A,B), geographic population variability (Fig. 3C), phonetic features (Figs. 4, 18), and acoustic categories (Fig. 5) are all captured by unsupervised latent space representations. Where possible, these distributional properties can (and should be) linked to specific abstract physical features of the signals, but our methods show that *a priori* feature-based compression is not a prerequisite to progress in understanding behaviorally relevant acoustic diversity. We used these latent projections to visualize sequential organization and abstract sequential models of song (Fig. 6) and demonstrated that in some cases latent approaches confer advantages over hand labeling or supervised learning (Fig. 7). We also projected vocalizations as continuous trajectories in latent space (Figs. 8, 9, 10, and 11). This provides a powerful method for studying sequential organization without discretizing vocal sequences [1]. In addition, we surveyed several deep neural network architectures (Fig. 12) that learn latent representations of vocal repertoires and systematically generate novel syllables from the features in latent space (Figs. 13, 14). Finally, we gave an example of how these methods can be combined in a behavioral experiment to study perception with psychometric precision, and in an acute electrophysiological experiment to understand representational encoding of parametrically varying natural vocal signals (Fig. 15).

### Latent and generative models in the biological sciences

Latent and generative models have shown increasing utility in the biological sciences over the past several years. As pattern recognition and representation algorithms improve, so will their utility in characterizing the complex patterns present in biological systems like animal communication. In neuroscience, latent models already play an important role in characterizing complex neural population dynamics [38]. Similarly, latent models are playing an increasingly important role in computational ethology [17], where characterizations of animal movements and behaviors have uncovered complex sequential organization [32, 89, 90]. In animal communication, pattern recognition using various machine learning techniques has been used to characterize vocalizations and label auditory objects [3, 27, 29, 33, 39, 66, 67]. Our work furthers this emerging research area by demonstrating the utility of unsupervised latent models for both systematically visualizing, characterizing, and generating animal vocalizations across a wide range of species.

### Discrete and continuous representations of vocalizations

Studies of animal communication classically rely on segmenting vocalizations into discrete temporal units. In many species, temporally segmenting vocalizations into discrete elements is a natural step in representing vocal data. In birdsong, for example, temporally distinct syllables are often well defined by clear pauses between highly stereotyped syllable categories. In many other species, however, vocal elements are either less clearly stereotyped or less temporally distinct, and methods for segmentation can vary based upon changes in a range of acoustic properties, similar sounds, or higher-order organization [1]. These constraints force experimenters to make decisions that can have profound effects on downstream analyses [29, 39]. We projected continuous latent representations of vocalizations ranging from highly stereotyped Bengalese finch song, to highly variable mouse USVs, and found that continuous latent projections effectively described useful aspects of spectro-temporal structure and sequential organization. In human speech, we found that continuous latent variable projections were able to capture sub-phoneme temporal dynamics that correspond to co-articulation. Collectively, our results show that continuous latent representations of vocalizations provide an alternative to discrete segment-based representations while remaining agnostic to segment boundaries, and without the need to segment vocalizations into discrete elements or symbolic categories. Of course, where elements can be clustered into clear and discrete element categories, it is important to do so. The link from temporally continuous vocalization to symbolically discrete sequences will be an important target for future investigations.

### Choosing a network architecture

The generative neural networks and machine learning models presented here are only a tiny sample of a very rapidly growing and changing field. We did not explore many of the potentially promising variants of adversarial networks or autoencoders (e.g. [73, 75, 91]) or any of the models that act directly on waveforms (e.g. [92]) or other classes of generative models (e.g. [93]). Each of these potentially possess different benefits in sampling from latent vocal spaces. Likewise, we did not rigorously explore differences between latent representations learned by different architectures of deep neural networks or the many other popular dimensionality reduction techniques like t-SNE [26]. The latent analyses presented here use UMAP, while in behavioral experiments we have been using VAEs [30], because they are computationally simple, tractable, easy to train, and enable generative output. The other network architectures we surveyed, as well as many emerging network architectures and algorithms, may offer promising avenues for generating even more realistic, higher fidelity vocal data, and for learning structure-rich latent feature spaces. Our brief survey is not meant to be exhaustive, but rather to serve as an introduction to many of the potentially rich uses of existing and future neural networks in generating and sampling from latent representational spaces of vocal data.

### Future work

The work presented here is a first step in exploring the potential power of latent and generative techniques in modeling animal communication. We touch only briefly on a number of questions that we find interesting and think important within the field of animal communication. Other researchers may certainly want to target other questions, and we hope that some of these techniques (and the provided code) may be adapted in that service. Our analyses were taken from a diverse range of animals, sampled in diverse conditions both in the wild and in the laboratory, and are thus not well controlled for variability between species. Certainly, as bioacoustic data becomes more open and readily available, testing large, cross-species, hypotheses will become more plausible. We introduced several areas in which latent models can act as a powerful tool to visually and quantitatively explore complex variation in vocal data. These methods are not restricted to bioacoustic data, however. Indeed many were designed originally for image processing. We hope that the work presented here will encourage a larger incorporation of latent and unsupervised modeling as a means to represent, understand, and experiment with animal communication signals in general. At present, our work exhibits the utility of latent modeling on a small sampling of the many directions that can be taken in the characterization of animal communication.

## 5 Methods

### 5.1 Datasets

The Buckeye [94] dataset of conversational English was used for human speech. The swamp sparrow dataset is from[18] and was acquired from [95]. The California thrasher dataset is from [6] and was acquired from BirdDB [96]. The Cassin’s vireo dataset is from [7] and was also acquired from BirdDB. The giant otter dataset was acquired from [97]. The canary song dataset is from [5] and was acquired via personal correspondence. Two zebra finch datasets were used. The first is a dataset comprised of a large number of motifs produced by several individuals from [98]. The second is a smaller library of vocalizations with more diverse vocalization types and a greater number of individuals than the motif dataset. It correspond to data from [99] and [100] and was acquired via personal correspondence. The white-rumped munia dataset is from [4]. The humpback whale dataset was acquired from Mobysound [101]. The house mice USV dataset was acquired from [65]. An additional higher SNR dataset of mouse USVs was sent from the same group via personal correspondence. The European starling dataset is from [3] and was acquired from [102]. The gibbon song is from [103]. The marmoset dataset was received via personal correspondence and was recorded similarly to [25]. The fruit bat data is from [104] and was acquired from [105]. The macaque data is from [23] and was acquired from [106]. The beaked whale dataset is from [51] and was acquired from [107]. The North American birds dataset is from [108] and was acquired from [50]. We used two Bengalese finch datasets. The first is from [8] and was acquired from [58]. The second is from [57].

#### Segmentation

Many datasets were made available with vocalizations already segmented either manually or algorithmically into units. When datasets were pre-segmented, we used the segment boundaries defined by the dataset authors. For all other datasets, we used a segmentation algorithm we call dynamic threshold segmentation (Fig. 16A). The goal of the algorithm is to segment vocalization waveforms into discrete elements (e.g. syllables) that are defined as regions of continuous vocalization surrounded by silent pauses. Because vocal data often sits atop background noise, the definition for silence versus vocal behavior was set as some threshold in the vocal envelope of the waveform. The purpose of the dynamic thresholding algorithm is to set that noise threshold dynamically based upon assumptions about the underlying signal, such as the expected length of a syllable or a period of silence. The algorithm first generates a spectrogram, thresholding power in the spectrogram below a set level to zero. It then generates a vocal envelope from the power of the spectrogram, which is the maximum power over the frequency components times the square root of the average power over the frequency components for each time bin over the spectrogram:

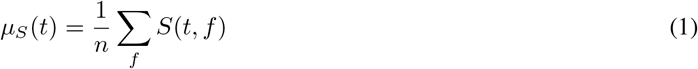

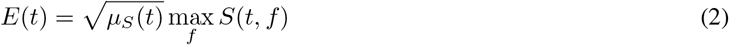

Where *E* is the envelope, *S* is the spectrogram, *t* is the time bin in the spectrogram, *f* is the frequency bin in the spectrogram, and *n* is the total number of frequency bins.

The lengths of each continuous period of putative silence and vocal behavior are then computed. If lengths of vocalizations and silences meet a set of thresholds (e.g. minimum length of silence and maximum length of continuous vocalization) the algorithm completes and returns the spectrogram and segment boundaries. If the expected thresholds are not met, the algorithm repeats, either until the waveform is determined to have too low of a signal to noise ratio and discarded, or until the conditions are met and the segment boundaries are returned. The code for this algorithm is available on Github [109].

#### Spectrogramming

Spectrograms are created by taking the absolute value of the one-sided short-time Fourier transformation of the Butterworth band-pass filtered waveform. Power is log-scaled and thresholded using the dynamic thresholding method described in the Segmentation section. Frequency ranges and scales are based upon the frequency ranges occupied by each dataset and species. Either frequency is logarithmically scaled over a frequency range using a Mel filter, or a frequency range is subsetted from the linearly frequency scaled spectrogram. Unless otherwise noted, all of the spectrograms we computed had a total of 32 frequency bins, scaled across frequency ranges relevant to vocalizations in the species.

To create a syllable spectrogram dataset (e.g. for projecting into Fig. 1), syllables are segmented from the vocalization spectrogram. To pad each syllable spectrogram to the same time length size, syllable spectrograms are log-rescaled in time then zero-padded to the length of the longest log-rescaled syllable.

#### Projections

Latent projections are either performed over discrete units (e.g. syllables) or as trajectories over continuously varying sequences. For discrete units, syllables are segmented from spectrograms of entire vocalizations, rescaled, and zero-padded to a uniform size (usually 32 frequency and 32 time components). These syllables are then projected either into UMAP or any of the several neural network architectures we used. Trajectories are projected from rolling windows taken over a spectrogram of the entire vocal sequence (e.g. a bout). The rolling window is a set length in milliseconds and each window is sampled as a single point to be projected into latent space. The window then rolls one frame (one time bin) at a time across the entire spectrogram, such that the number of samples in a bout trajectory is equal to the number of time frames in the spectrogram. These time bins are then projected into UMAP latent space.

#### Neural networks

Neural networks were designed and trained in Tensorflow 2.0 [110]. We used a combination of convolutional layers and fully connected layers for each of the network architectures except the seq2seq network, which is also comprised of LSTM layers. We generally used default parameters and optimizers for each network, for example, the ADAM optimizer and rectified linear (ReLU) activation functions. Specific architectural details can be found in the GitHub repository.

#### Clusterability

We used the Hopkin’s statistic [40] as a measure of the clusterability of datasets in UMAP space. In our case, the Hopkin’s statistic was preferable over other metrics for determining clusterability, such as the Silhouette score [111] because the Hopkin’s statistic does not require labeled datasets or make any assumptions about what cluster a datapoint should belong to. The Hopkin’s statistic is part of at least one birdsong analysis toolkit [95].

The Hopkin’s statistic compares the distance between nearest neighbors in a dataset (e.g. syllables projected into UMAP), to the distance between points from a randomly sampled dataset and their nearest neighbors. The statistic computes clusterability based upon the assumption that if the real dataset is more clustered than the randomly sampled dataset, points will be closer together than in the randomly sampled dataset. The Hopkin’s statistic is computed over a set *X* of *n* data points (e.g. latent projections of syllables of birdsong), where the set *X* is compared with a baseline set *Y* of *m* data points sampled from either a uniform or Gaussian distribution. We chose to sample *Y* from a uniform distribution over the convex subspace of *X*. The Hopkin’s metric is then computed as:

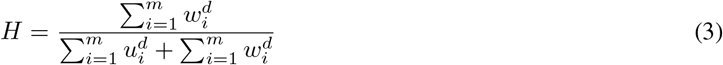

Where *u*_*i*_ is the distance of *y*_*i*_ ∈ Y from its nearest neighbor in *X* and *w*_*i*_ is the distance of *x*_*i*_ ∈ X from its nearest neighbor in *X*. Thus if the real dataset is more clustered than the sampled dataset, the Hopkin’s statistic will approach 0, and if the dataset is less clustered than the randomly sampled dataset, the Hopkin’s statistic will sit near 0.5. Note that the Hopkin’s statistic is also commonly computed with 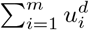 in the numerator rather than 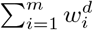, where Hopkin’s statistics closer to 1 would be higher clusterability, and closer to 0.5 would be closer to chance. We chose the former method because the range of Hopkin’s statistics across datasets were more easily visible when log transformed.

#### Comparing algorithmic and hand-transcriptions

Several different metrics can be used to measure the overlap between two separate labeling schemes. We used four metrics that capture different aspects of similarity to compare hand labeling to algorithmic clustering methods ([60]; Table 1). Adjusted Mutual Information is an information-theoretic measure that quantifies the agreement between the two sets of labels, normalized against chance. Completeness measures the extent to which members belonging to the same class (hand label) fall into the same cluster (algorithmic label). Homogeneity measures whether all clusters fall into the same class in the labeled dataset. V-Measure is the harmonic mean between homogeneity and completeness. We found that HDBSCAN and UMAP showed higher similarity to human labeling than k-means in nearly all metrics across all three datasets.

## Data Availability

All of the vocalization datasets used in this study were acquired from external sources, most of them hosted publicly online. The behavioral and neural data are part of a larger project and will be released alongside that manuscript.

## Code Availability

The python code written specifically for this paper is available at Github.com/timsainb/AVGN_paper. A cleaner and more maintained code base is additionally available at Github.com/timsainb/AVGN.

## Acknowledgments

Work supported by NSF GRF 2017216247 and an Annette Merle-Smith Fellowship to T.S. and NIH DC0164081 and DC018055 to T.Q.G. We additionally would like to thank Kyle McDonald and his colleagues for motivating some of our visualization techniques with their work on humpback whale song.

## Ethics statement

Procedures and methods comply with all relevant ethical regulations for animal testing and research and were carried out in accordance with the guidelines of the Institutional Animal Care and Use Committee at the University of California, San Diego (S05383).

## Notes

#### Summary of Updates

Fixed error in figure capture where Fig 3A and Fig 3B were reversed.

https://github.com/timsainb/avgn_paper

